# Locomotion-invariant prefrontal–thalamic goal states organize spatially aligned episode-specific hippocampal maps

**DOI:** 10.64898/2026.04.09.716651

**Authors:** Zahra Golipour, Shao-Fan Yen, Cansu Üstüner, Hiroshi T. Ito

**Affiliations:** University of Lausanne, Switzerland; Max Planck Institute for Brain Research, Germany

## Abstract

Animals repeatedly traverse the same environment to pursue different goals, yet the hippocampus must preserve a stable spatial map while keeping individual episodes distinct. Here we show that, when animals navigate the same maze under different goal configurations, hippocampal CA1 segregates navigation episodes by encoding goal state along a population dimension orthogonal to the spatial coding subspace, rather than by reorganizing spatial representations themselves, allowing episode-specific maps to remain spatially aligned. This goal-state signal is supplied by a prefrontal–thalamic pathway, in which population activity in medial prefrontal cortex and nucleus reuniens forms persistent representations across locomotion and immobility and is reliably reinstated when previously experienced goal configurations recur. Silencing the nucleus reuniens selectively abolishes CA1 goal-state coding by disrupting goal-axis separation and goal-biased pre-navigation spike sequences while sparing spatial coding. Together, these findings identify a circuit- and population-level mechanism that enables episode-specific hippocampal representations to coexist within spatially consistent maps independent of locomotor state, linking internal goal states to navigation and planning.

## Main

A central challenge for memory systems is to represent distinct experiences that occur within the same physical environment^1–6^. The hippocampus is well known for its spatial coding properties^2,7,8^, yet spatially similar or even identical trajectories can correspond to distinct episodes. How the hippocampal network preserves a stable spatial map while simultaneously maintaining episode-specific representations therefore remains unclear. Classical accounts emphasize activity changes driven by sensory, motor, or contextual modulation^9–23^, but such mechanisms typically alter spatial representations themselves, making it unclear how multiple episodes can coexist within the same environment without mutual interference.

Recent work has shown that hippocampal population activity can incorporate non-spatial variables, including internal states, task demands, and latent cognitive dimensions^24,25^. These findings raise the possibility that the hippocampus may separate episodes by allocating them to distinct subspaces of its population activity. However, it remains unknown whether a dedicated population dimension exists that can partition episodes while preserving the integrity of spatial coding subspaces, and, critically, which upstream circuits supply the internal contextual signals required to drive such partitioning. Identifying this mechanism is essential for understanding how the hippocampus maintains stable maps while flexibly supporting planning, decision-making, and episodic recall^26–31^.

An additional problem in linking hippocampal function in spatial navigation and memory concerns the locomotion dependence of hippocampal activity. Hippocampal neural dynamics differ profoundly between locomotion and immobility^32–35^, yet episodic states must be maintained irrespective of movement. The mechanisms that support such locomotion-invariant state representations in hippocampal population activity have remained elusive.

Prefrontal–thalamic circuits are strong candidates for generating these internally defined contextual states^36–40^. The medial prefrontal cortex (mPFC) and the thalamic nucleus reuniens (NR) form a pathway that regulates hippocampal states and is implicated in working memory, behavioral flexibility, and goal-directed planning^17,41–44^. However, whether this pathway provides the hippocampus with a stable and reinstatable goal-state representation—capable of partitioning episodes independently of ongoing behavior—remains unknown. Such a mechanism would offer a principled solution to the interference problem, allowing multiple episode-specific yet spatially-consistent hippocampal maps to coexist within the same environment.

These considerations motivate the hypothesis that goal-state information from the mPFC–NR pathway is integrated into the hippocampal population code along a neural dimension orthogonal to the spatial coding subspace, enabling the segregation of episode-specific maps without altering spatial structure. Moreover, if this externally supplied state signal is independent of locomotor state, it could enable the hippocampus to maintain a locomotion-invariant neural state that is dissociable from spatial position coding.

To test this hypothesis, we developed a goal-directed navigation task in a linear maze in which rats traverse identical spatial trajectories while reward locations alternate across blocks. This design isolates the impact of goal-state changes on neural dynamics by ensuring that spatial and sensory inputs remain constant—an approach that is difficult to achieve in conventional two-dimensional maze environments. Using large-scale electrophysiological recordings from mPFC, NR, and hippocampal CA1, combined with optogenetic perturbations of NR neurons, we examine how goal-related information is causally transmitted to the hippocampus and incorporated into its population coding structure. Crucially, we further assess whether this goal-state code is expressed across both locomotion and immobility, thereby unifying episode-specific representations across locomotor states. This framework enables us to identify the circuit-level mechanism that allows spatially aligned yet episode-specific hippocampal maps to coexist and to determine how such maps are selected during planning and navigation.

## Results

### CA1 neurons maintain stable spatial codes across journeys targeting different goals

To examine how navigational goals shape hippocampal activity, we recorded CA1 neurons by implanting a microdrive carrying 28 independently movable tetrodes bilaterally in dorsal CA1 while rats performed a goal-directed navigation task on a linear maze equipped with ten equidistant water-delivery wells (1,861 neurons from 9 rats, 207 ± 13 neurons per animal per session; Figs. 1a, 1b, Extended Data Figs 1 and 2). Throughout the task, rats navigated a sequence of three goal wells—two fixed at the trackends and a middle goal that changed after a random number of trials (4–15). Each trial comprised four journeys between the immediately adjacent goal wells, encompassing both outbound and inbound runs. A block with new goals began with an introduction trial in which water was pre-delivered at the upcoming goal well, and LEDs under all ten wells were illuminated to signal the goal change. After this initial pre-water trial, rats were required to lick the designated goal well for a fixed duration (0.8–1.5 s) before receiving water, ensuring that correct performance depended on estimating the goal location.

**Figure 1:**
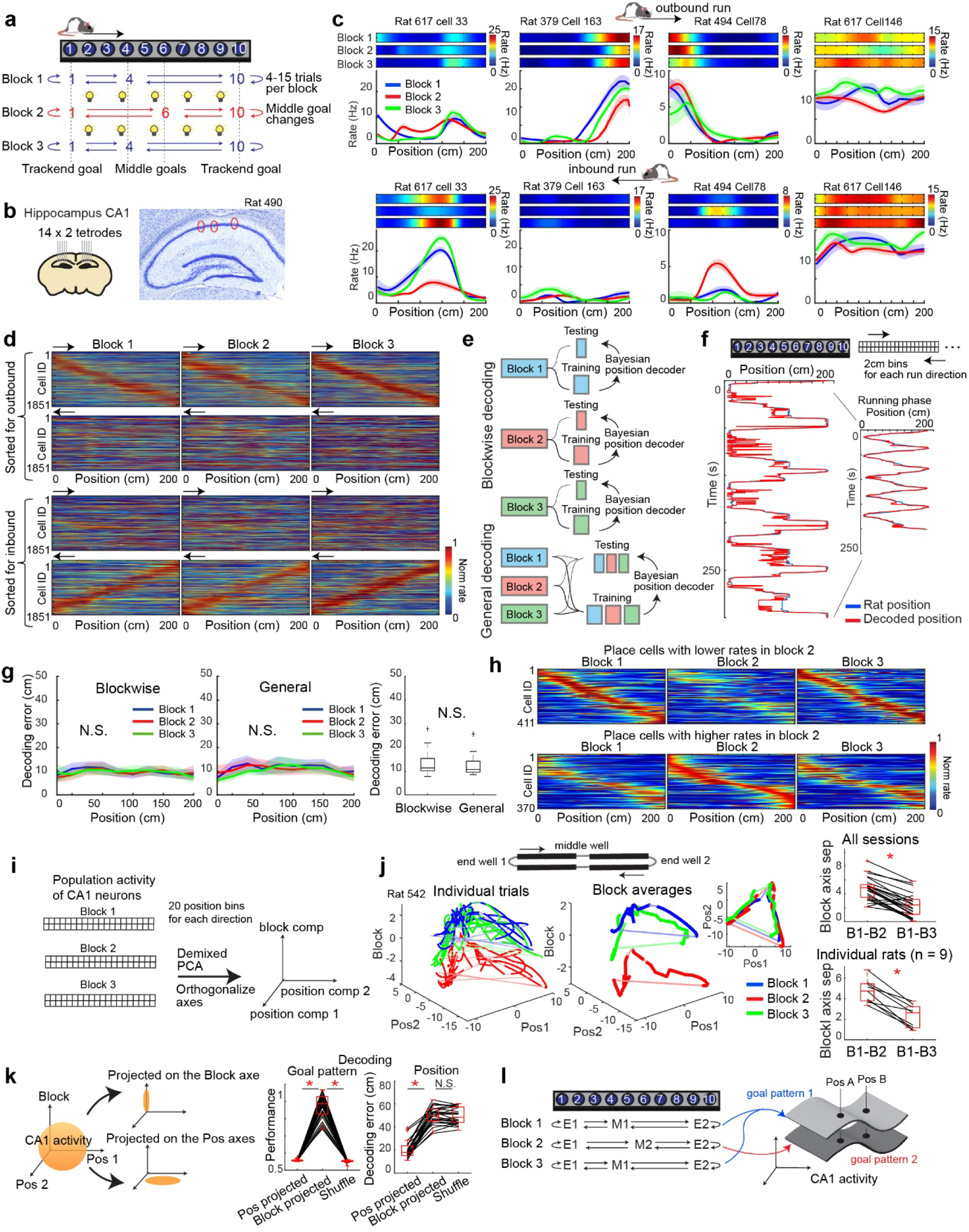
CA1 segregates distinct goal-directed navigation episodes into spatially aligned maps. **a,** Schematic of the navigation task, illustrating the maze design and task rules. The animal sequentially navigates three goal locations, with one complete cycle comprising four goal-directed journey segments. Each trial block consists of a random number of trials (4–15). The identity of the middle goal changes between blocks. Animals performed the task at >80% accuracy and adapted to goal changes within 1–2 cycles (Extended Data Fig. 3). **b,** A microdrive carrying 28 tetrodes was implanted bilaterally in hippocampal CA1. The right panel shows a Nissl-stained coronal section with red circles indicating tetrode tracks. **c,** Firing-rate maps of four representative CA1 neurons across three trial blocks, shown separately for each running direction in the linear maze. **d,** Color-coded firing-rate maps of all CA1 neurons recorded in individual blocks for each running direction. Neurons are sorted by the location of their firing fields along the maze: the top two panels sorted for outbound journeys and the bottom two for inbound journeys. **e,** Schematic of the Bayesian position-decoding strategy based on CA1 ensemble activity. For blockwise decoding, training and testing were performed within individual blocks. For general decoding, data from all blocks were pooled and randomly interleaved for decoder training and testing. **f,** Example traces of the animal’s actual position and the decoder’s estimated position, illustrating accurate decoding, particularly during the running phase. **g,** Decoding errors as a function of maze position across trial blocks. The box plot on the right compares the mean decoding errors of the two decoders and shows no significant difference. p > 0.05, Wilcoxon signed-rank test. **h,** Color-coded firing-rate maps of all CA1 place cells, plotted separately for neurons with lower or higher firing rates in block 2 relative to blocks 1 and 3. **i,** Schematic illustrating the demixed PCA approach used to extract two position components and one block component, followed by orthogonalization of the axes. **j,** Three-dimensional projections of CA1 population activity onto the block component and two position components. Left, activity from individual trials; right, block-averaged activity. The upper-right inset shows activity traces projected onto the two position components. Activity from block 2 is separated from blocks 1 and 3 along the block axis, while overlapping along the position axes. The two box plots on the left quantify block-axis separation between blocks 1 and 2 versus blocks 1 and 3, either across 25 sessions (*p < 0.001) or from 9 individual rats (*p = 0.008). Wilcoxon signed-rank test. **k,** Assessment of information content along each projection axis. CA1 population activity was projected onto either the block component or the position components, and decoding performance was evaluated accordingly (right two box plots). Goal-related and spatial information occupy separate projection axes in the population activity. *p < 0.05, Wilcoxon signed-rank test. **l,** Schematic illustrating the orthogonal coding scheme in CA1 population activity. Depending on the goal configuration, CA1 forms two spatially aligned maps that preserve spatial coding consistency while segregating distinct navigation episodes targeting different goals.

**Figure 2:**
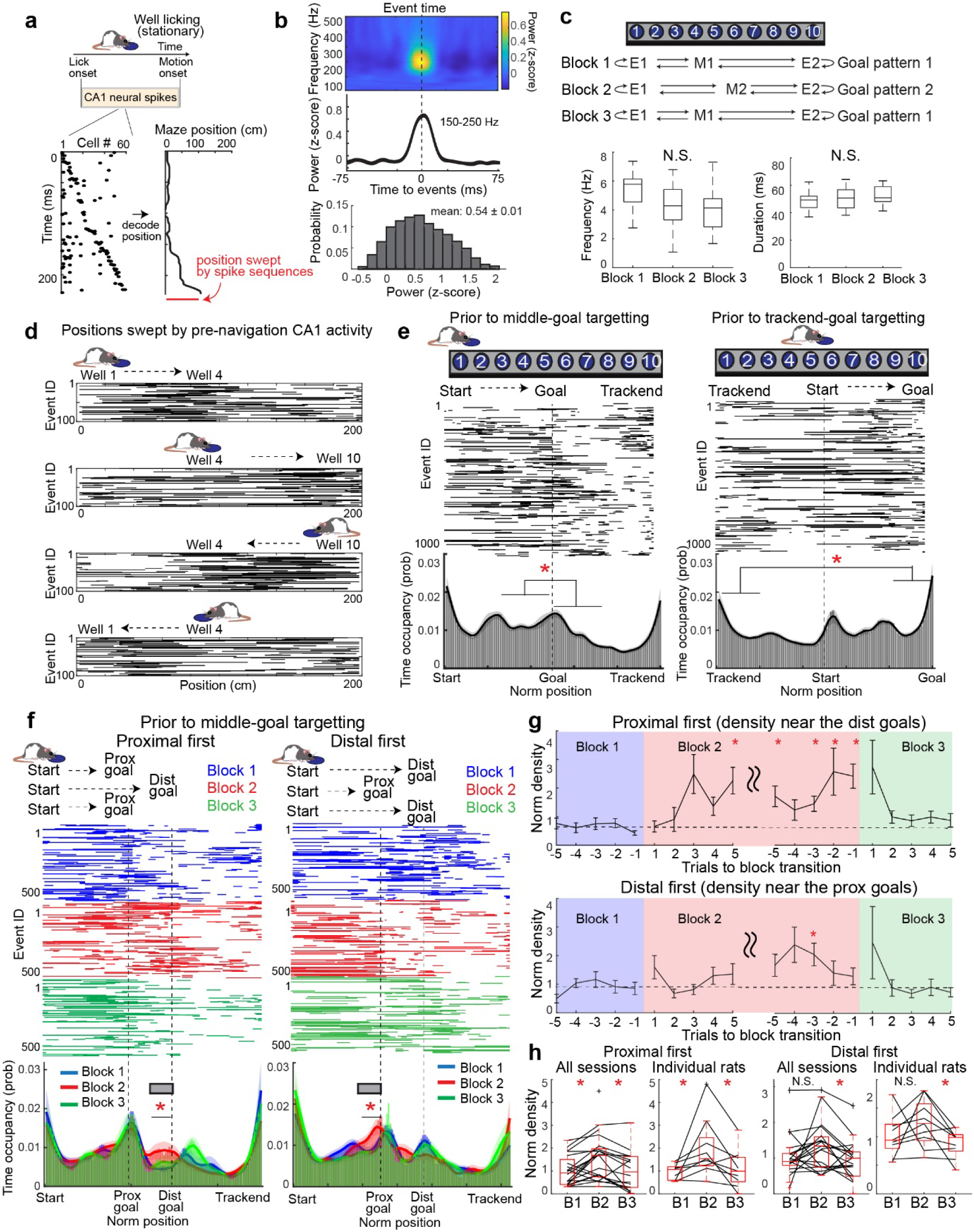
Pre-navigation CA1 spike sequences flexibly adapt to upcoming goal locations. **(A)** Schematic showing detection of pre-navigation spike sequences and definition of corresponding swept positions. **b,** Plots showing the spectral power in CA1 LFPs averaged around the times of detected spike sequence events. Top: color-coded spectral power; middle: average power at 150 –250 Hz frequency range; bottom: histogram showing the probability of z-scored power of individual spike sequence events. **c,** Comparison of frequency and duration of detected spike sequence events across blocks. p > 0.05, Kruskal-Wallis test. **d,** Spatial distributions of individual spike-sequence events along maze positions at different trial phases during quiescence periods before movement starts. **e,** Spatial distributions (top) and time-occupancy probabilities (bottom) of spike sequences across maze positions, shown separately for periods preceding navigation toward middle goals (right) and toward trackend goals (left). All plots are normalized to the goal and trackend positions, respectively. Left: The ratio of occupancy at the middle goal relative to the first quadrant is significantly larger than that of the third quadrant relative to the middle goal. *p < 0.001 in Wilcoxon rank-sum. Right: Time-occupancy probabilities are significantly higher in the final quadrant (near the upcoming goal) than in the first quadrant (near the previous goal). *p = 0.006, Wilcoxon rank-sum test. **f,** Spatial distributions (top) and time-occupancy probabilities (bottom) of pre-navigation spike sequences across maze positions. Left: Trials in which animals targeted a proximal goal in blocks 1 and 3 and a distal goal in block 2. Right: Trials in which animals targeted a distal goal in blocks 1 and 3 and a proximal goal in block 2. Black lines indicate regions with significantly higher occupancy in block 2 compared to blocks 1 and 3. *p < 0.05 in Wilcoxon rank-sum test. **g,** Trial-by-trial analysis of time-occupancy probability within the region of interest defined in **f**. *p < 0.05, Wilcoxon rank-sum test relative to block 1. **h,** Box plots summarizing time-occupancy probabilities within the region of interest across all 25 sessions from 9 rats. Some data points are missing due to the absence of detected sequence events in specific blocks. *p < 0.05, Wilcoxon signed-rank test.

Each recording session consisted of three consecutive blocks. Blocks 1 and 3 shared the same goal configuration, whereas block 2 featured a different middle goal. For each session, the middle goals were randomly selected from wells 4–7. All wells were identical in shape, and behavioral sessions were conducted under minimal lighting to ensure that navigation was not directly guided by specific sensory cues. Overall, the animals performed the task with high accuracy (86.0 ± 1.8%) and exhibited smooth goal-targeting without slowing in the middle of journeys (Extended Data Fig. 3).

We first assessed the spatial tuning of individual CA1 neurons as rats targeted different middle-goal wells in the maze. We found that the majority of place cells maintained consistent spatial tuning despite changes in goal location, although they exhibited pronounced differences between the two running directions (Figs. 1c and 1d). Consistent with this directional dependence, a Bayesian decoder of the animal’s position^26,45^ trained on one running direction performed significantly worse when tested on the opposite direction (Extended Data Fig. 3), in line with previous studies demonstrating largely independent spatial codes between running directions^46,47^. Given this directional specificity, we quantified the proportions of spatially and non-spatially tuned neurons separately for each running direction (Extended Data Fig. 3).

Among spatially tuned CA1 neurons (spatial information > 0.25 bits/spike), 31.8% did not maintain stable firing between the identical-goal blocks (blocks 1 and 3), and 9.3% exhibited global remapping in response to changes in goal location. However, the majority of spatial neurons (58.9%) maintained stable spatial fields across trial blocks despite changes in the middle-goal position.

To further validate this observation at the population level, we tested whether the animal’s position could be decoded consistently from CA1 ensemble activity across trial blocks. We applied a Bayesian decoder trained and tested on CA1 population activity (Figs. 1e, 1f and Extended Data Fig. 3). To assess whether CA1 neurons preserve accurate position coding despite goal changes, we compared position estimation errors from decoders trained separately on each block (“blockwise decoder”) with those from a decoder trained across blocks with different goal configurations (“general decoder”). Because spatial coding differs between the two running directions, we constructed separate decoding templates for each direction and applied them to each time bin, yielding estimates of both the animal’s position and running direction (Extended Data Fig. 3).

We found that the blockwise and general decoders exhibited comparable performance across individual blocks, with no significant difference in decoding accuracy (Fig. 1g; decoding errors: blockwise, 12.7 ± 0.9 cm; general, 12.6 ± 0.8 cm; p = 0.677, Wilcoxon signed-rank test). These results confirm that CA1 spatial coding remains stable across journeys targeting different goals within the same environment.

### CA1 forms spatially aligned maps that orthogonally separate task blocks associated with different goals

The observation of stable position decoding does not imply that CA1 neuronal activity is insensitive to changes in goal location. Indeed, we observed that even place cells with stable spatial tuning exhibited systematic changes in peak firing rates, either increases or decreases, during block 2 compared to blocks 1 and 3 (Fig. 1h), suggesting that these rate differences may reflect changes in middle-goal location. Similar modulations of place-cell activity have previously been described as “rate remapping,” in which firing rates change according to sensory features of the environment while spatial tuning is preserved^10^.

To further characterize this effect, we quantified the remapping properties of CA1 neurons. Among neurons that exhibited consistent spatial tuning across blocks, 40.2% showed significant rate changes within their firing fields corresponding to differences in goal location (Extended Data Fig. 3). Comparable goal-dependent rate changes were also observed in 32.0% of non-spatial cells (spatial information < 0.25 bits/spike; Extended Data Fig. 3). To evaluate the impact of rate remapping on population-level activity, we compared population vector (PV) correlations of ensemble firing patterns for spatial and non-spatial cells. PV correlations were significantly lower between blocks with different goals (blocks 1 and 2) than between blocks with the same goals (blocks 1 and 3) for both spatial and non-spatial cells (spatial cells: blocks 1–2, 0.614 ± 0.028; blocks 1–3, 0.774 ± 0.018; non-spatial cells: blocks 1–2, 0.232 ± 0.020; blocks 1–3, 0.337 ± 0.033; p < 0.001 for both cell types; Friedman test; Extended Data Fig. 3). These results indicate that differences in goal distributions are reflected in CA1 ensemble activity through a rate-remapping mechanism.

These observations at the level of individual neurons suggest that CA1 populations can represent goal-related patterns independently of spatial representations, thereby preserving stable spatial coding while encoding distinct goal states. To test this idea directly, we applied demixed principal component analysis (dPCA)^48^, a linear dimensionality-reduction method that separates population activity into components associated with predefined task parameters (Fig. 1i). This analysis decomposed CA1 ensemble firing into three task-relevant components: a block component corresponding to individual task blocks with distinct goal configurations, and two position components capturing the animal’s position along the maze in each of the two running directions. Together, these components reveal a low-dimensional population structure associated with block and position variables. The block component accounted for 4.9 ± 0.4% of the variance, whereas the two position components together explained 40.1 ± 1.5%. Although dPCA does not enforce orthogonality among components, we verified that these components were nearly orthogonal (Extended Data Fig. 3). We further applied a Gram–Schmidt procedure to ensure orthogonality of the three components for visualization (see Methods).

Projecting CA1 population activity onto these components revealed that positional information was captured by the two position axes, whereas differences between goal patterns were expressed along the block axis (Fig. 1j). The separation along the block axis between blocks 1 and 2, which involved different middle-goal locations, was significantly larger than that between blocks 1 and 3, which shared the same goal configuration (blocks 1–2: 3.73 ± 0.29; blocks 1–3: 1.91 ± 0.28 arbitrary units; p < 0.001 across sessions).

To further assess the segregation of spatial and goal information, we projected CA1 population activity onto either the block axis or the position axes and quantified how much goal-pattern and positional information could be recovered using linear discriminant analysis (LDA; Fig. 1k). Goal information was selectively preserved when activity was projected onto the block axis but dropped to near chance when projected onto the position axes. Conversely, accurate position decoding was achieved when decoding was performed along the position axes but deteriorated when decoding was performed along the block axis. These results confirm that spatial position and goal pattern are encoded in largely orthogonal subspaces of the CA1 population activity manifold. We obtained the same conclusion using an alternative decoding approach that did not rely on dPCA (Extended Data Fig. 3).

Together, these findings indicate that CA1 employs an orthogonal coding scheme that segregates goal distributions while preserving stable spatial representations within the same environment, consistent with the formation of spatially aligned parallel maps (Fig. 1l).

### CA1 spike sequences prior to navigation distinguish upcoming destinations

If goal-state representations truly function as episode selectors rather than transient task signals, they should be expressed not only during active navigation but also in the internally generated hippocampal dynamics that precede movement. While the results above demonstrate that CA1 population activity distinguishes goal patterns during active navigation, it remains unclear whether CA1 also expresses distinct activity dynamics during the awake quiescent period preceding navigation onset. Hippocampal neurons are known to generate brief spike sequences that recapitulate future or past trajectories ^26–29^, but previous studies have largely examined fixed-goal tasks or tasks in which different goals are associated with distinct navigation trajectories. It has therefore remained unclear whether and how goal differences directly influence pre-navigation internal dynamics in the hippocampus.

To examine the spatial representations embedded in CA1 pre-navigation activity, we applied the same Bayesian position decoder trained on population activity during active locomotion to neural activity recorded during the 3-s period immediately preceding navigation onset, when animals were stationary at the start wells (Fig. 2a). Consistent with previous studies^26,45^, we used short decoding windows to capture time-compressed dynamics (20-ms bins shifted every 5 ms; see Methods). Population activity was classified as a pre-navigation sequence event if it met predefined criteria for duration (>20 ms), trajectory smoothness (position jumps between bins <20 cm), and spatial extent (> 25 cm from the animal’s actual position). As previously reported^26,28,29^, these events were predominantly accompanied by sharp-wave ripples in the CA1 local field potential (Fig. 2b).

We first examined the spatial coverage of detected pre-navigation sequence events. The distribution of represented positions differed markedly depending on the animal’s current location and its upcoming destination (Fig. 2d), indicating that pre-navigation CA1 activity is dynamically modulated by navigational context. Prior to journeys targeting the middle goals, pre-navigation sequences preferentially represented positions spanning from the start well to the middle-goal location, with a sharp decline beyond the goal (Figs. 2e and Extended Data Fig. 3). In contrast, prior to journeys targeting the trackend goals, sequence activity showed a modest but significant bias toward future goal locations relative to recently traversed paths (Figs. 2e and Extended Data Fig. 3).

We further examined whether these dynamics differed between running directions as the Bayesian decoder can infer the animal’s running direction from the direction-selective firing of place cells. For the middle-goal–targeting journeys, no significant directional bias was detected. However, prior to trackend–goal–targeting journeys, a predictive bias toward the future goal emerged selectively when decoding was aligned with the animal’s upcoming movement direction (Extended Data Fig. 3), confirming the predictive nature of these pre-navigation sequences.

Finally, we examined how pre-navigation sequences adapted to changes in middle-goal location across trial blocks. Sequence density increased near the newly assigned goal location in block 2 compared to block 1 and decreased again when the original goal configuration was reinstated in block 3 (Figs. 2f–h). These effects were consistent across sessions and animals. The fact that this goal-dependent modulation emerges even when the animal is located at the exact same positions suggests that pre-navigation CA1 activity reflects goal-specific internal dynamics rather than differences in spatial or sensory input.

### Neural activity in mPFC and NR segregates task blocks associated with distinct goals

Our results thus far show that CA1 population activity encodes both spatial position and goal patterns along distinct coding dimensions. This organization suggests that goal-related information may be provided to the hippocampus by upstream regions. While the medial entorhinal cortex is a major source of spatial information to the hippocampus^8,49^, it remains unclear how CA1 receives information about differences in goal distributions during navigation. Based on prior evidence that trajectory-related signals are transmitted through the medial prefrontal cortex (mPFC)–nucleus reuniens (NR)–CA1 pathway^17^, we hypothesized that goal-related information is conveyed to CA1 via this circuit.

To test this hypothesis, we recorded neuronal population activity from mPFC and NR using Neuropixels probes while rats performed the same navigation task (mPFC: 708 neurons from 3 animals; NR: 503 neurons from 3 animals; Figs. 3a, 3b, and Extended Data Fig. 4). We first assessed whether information about the upcoming goal could be decoded from population activity at navigation onset. Decoding accuracy remained at chance levels in both regions (Extended Data Fig. 5), indicating that neither mPFC nor NR robustly encodes the immediate next goal of individual journeys, in contrast to the prospective goal coding reported previously in orbitofrontal cortex^40^.

**Figure 3:**
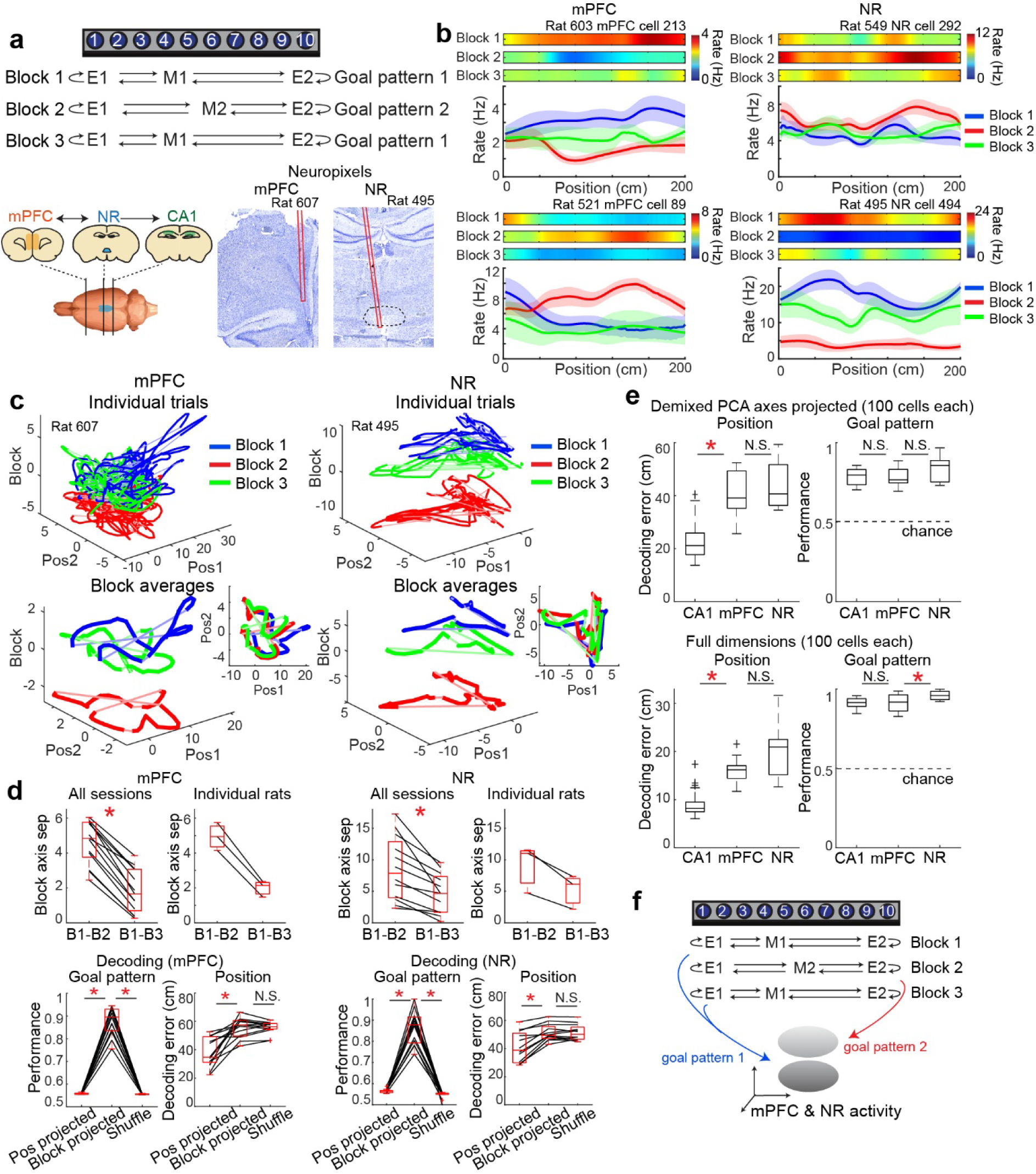
Activity in mPFC and NR segregates task blocks associated with different goals. **a,** Top: Task structure comprising three blocks with changes in the middle-goal location. Bottom: Schematic of the mPFC–NR–CA1 circuit and Nissl-stained coronal sections showing the tracks of Neuropixels 1.0 probes in mPFC and NR. **b,** Example firing patterns of two representative neurons from mPFC and two from NR. **c,** Three-dimensional projections of mPFC and NR population activity onto the block component and two position components, as in Fig. 1j. Top: Activity from individual trials. Bottom: Block-averaged activity. The upper-right inset shows activity projected onto the two position components. Activity from block 2 is separated from blocks 1 and 3 along the block axis, whereas activity along the position axes shows little consistency across blocks. **d,** Top: Box plots quantifying separation along the block axis between blocks 1 and 2 versus blocks 1 and 3, computed across 13 sessions (mPFC), 12 sessions (NR), and from 3 individual rats. Separation is significant in both regions. *p < 0.05, Wilcoxon signed-rank test. Bottom: Assessment of information content along each projection axis. Population activity was projected onto either the block component or the position components, and decoding performance was evaluated accordingly. Goal-related and spatial information are segregated into distinct projection axes of the population activity. *p < 0.05, Wilcoxon signed-rank test. **e,** Comparison of information content across CA1, mPFC, and NR using matched numbers of neurons across regions (100 cells). Decoding performance was assessed either on dPCA-derived orthogonalized axes (top) or using full-dimensional LDA decoding (bottom). Goal-pattern decoding performance was comparable across regions, whereas spatial-position decoding was significantly higher in CA1. *p < 0.05, Wilcoxon rank-sum test. **f,** Schematic illustrating the proposed neural representational structure in mPFC and NR. These regions robustly segregate goal patterns but exhibit comparatively weak spatial-position coding, consistent with a role in state coding rather than in forming spatial maps, unlike CA1.

We next examined whether population activity in mPFC and NR segregates task blocks associated with different goal patterns, analogous to the segregation observed in CA1. Applying demixed principal component analysis (dPCA) to the population activity, we decomposed neural responses into block-related and position-related components (explained variance: goal, mPFC 7.2 ± 0.9%, NR 19.6 ± 2.5%; position, mPFC 56.5 ± 5.3%, NR 30.1 ± 3.4%; Fig. 3c and Extended Data Fig. 6). After orthogonalizing the extracted components, we visualized population trajectories in the low-dimensional space defined by the block and position axes (Fig. 3c). In both mPFC and NR, trajectories corresponding to blocks with different goal patterns (blocks 1 and 2) were more strongly separated along the block axis than trajectories corresponding to blocks with identical goal patterns (blocks 1 and 3). Quantitatively, separation along the block axis was significantly larger between blocks 1 and 2 than between blocks 1 and 3 in both regions (mPFC: 4.64 ± 0.34 versus 1.82 ± 0.34; NR: 8.61 ± 1.48 versus 4.67 ± 0.91; p < 0.001, Wilcoxon signed-rank test). However, in contrast to CA1, population trajectories in mPFC and NR exhibited less consistent structure along the position axes (Fig. 3c, insets).

To further quantify the information carried along these axes, we projected population activity onto either the block component or the position components and assessed decoding performance using linear discriminant analysis (Fig. 3d). In both mPFC and NR, goal-pattern decoding performance remained high when decoding was performed along the block axis (goal classification performance: mPFC 0.88 ± 0.02, NR 0.86 ± 0.02) but dropped to near chance when activity was projected onto the position axes. Although position information could be decoded above chance from the position axes, overall position decoding accuracy in mPFC and NR was substantially lower than that observed in CA1 (decoding errors: mPFC 38.4 ± 2.9 cm, NR 40.9 ± 3.1 cm; Fig. 3e), indicating weaker spatial fidelity in these regions.

To directly compare coding properties across regions, we subsampled neural populations to matched sizes (100 neurons per session). Under these conditions, position decoding accuracy remained significantly higher in CA1 than in mPFC or NR, whereas goal-pattern decoding performance was comparable across all three regions, regardless of whether decoding was performed on dPCA-projected activity or on full-dimensional population activity (Fig. 3e).

Together, these results indicate that, unlike CA1 which encodes both spatial and goal-state information along largely independent dimensions, population activity in mPFC and NR primarily functions to segregate navigation episodes associated with different goal patterns, with relatively limited representation of spatial position. This pattern is consistent with the idea that the mPFC–NR pathway conveys a goal-state or contextual signal to the hippocampus, which is then integrated with spatial information in CA1 to form spatially aligned goal-specific population representations.

### Goal states are persistently represented across mPFC, NR, and CA1 during both locomotion and immobility

The analyses above demonstrate that mPFC and NR carry goal-state information during active navigation. A prominent feature of CA1 activity, however, is the presence of goal-associated spike sequences that occur during periods of immobility (Fig. 2). This raises the question of whether goal-state representations in the mPFC–NR–CA1 circuit extend beyond locomotion and persist during stationary behavioral states.

To address this question, we applied linear discriminant analysis (LDA) to classify blockwise goal patterns across entire recording sessions using time-segmented population activity, irrespective of the animal’s behavioral state (Fig. 4a). The decoder was trained on mean firing rates of neural ensembles computed from randomly sampled 2-s segments (equal numbers of segments from blocks 1 and 3, and twice as many from block 2 to balance class labels) and tested on other 2-s segments withheld from training (see Methods). Importantly, training segments were selected without regard to the animal’s position, movement, or locomotor state.

**Figure 4:**
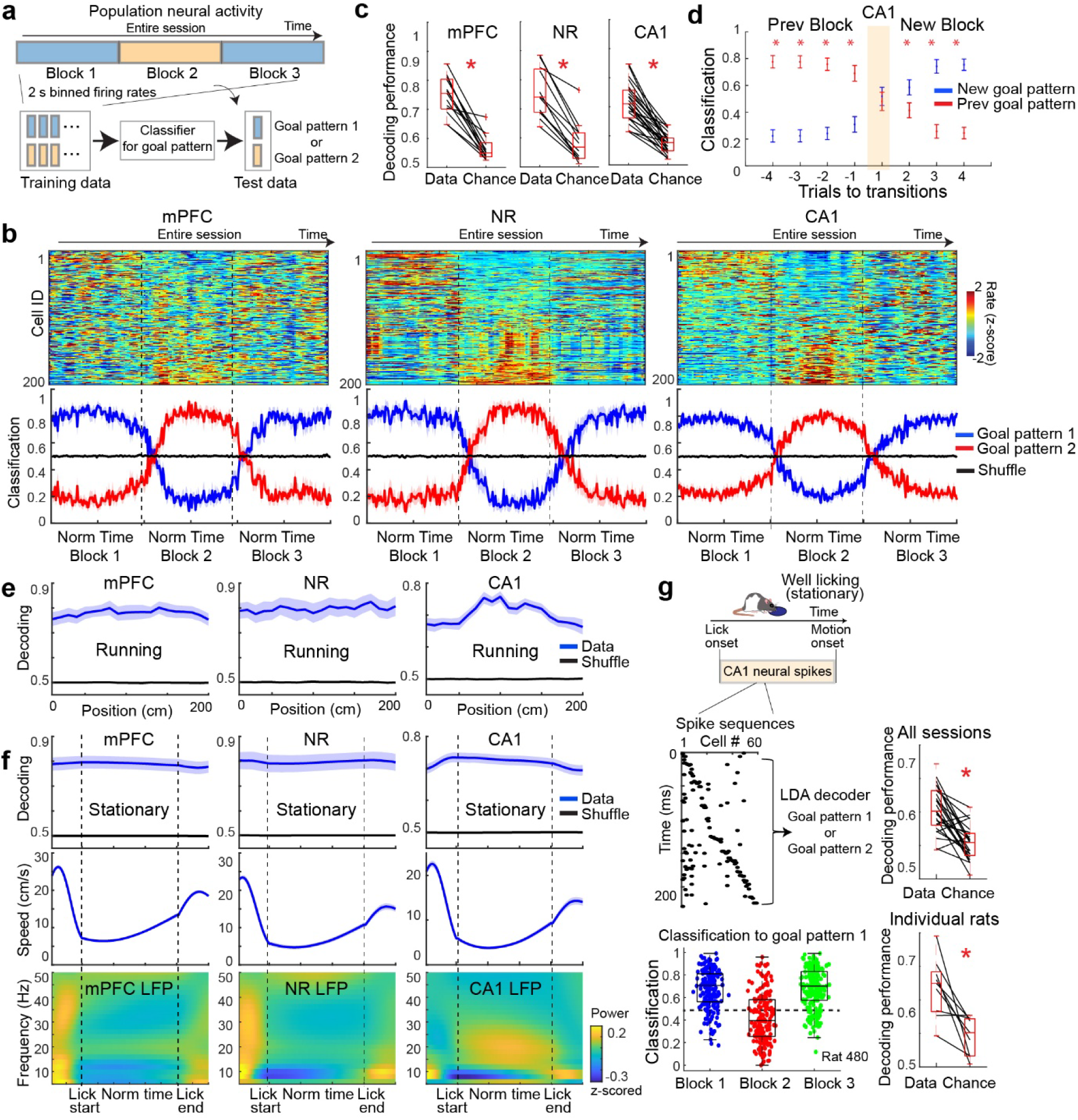
Persistent goal-state coding across the mPFC-NR-CA1 circuit throughout running and stationary states. **a,** Schematic illustrating the decoding strategy for goal-pattern discrimination. Ensemble neural activity was segmented into 2-s bins. For training, 15 bins were randomly sampled from blocks 1 and 3 and 30 bins from block 2, and the resulting classifier was used to predict the goal pattern of withheld test bins. **b,** Top: Color-coded, z-scored neural activity of 200 randomly sampled neurons across the entire recording session spanning blocks 1–3. Neurons are sorted by the magnitude of their firing-rate differences between goal patterns 1 and 2. Time axes are normalized to the duration of each block. Bottom: Goal-pattern classification probabilities plotted on the same time axis. **c,** Box plots showing mean goal-pattern decoding performance from neural ensembles in mPFC, NR, and CA1, compared with chance-level performance estimated by shuffled labels (Extended Data Fig. 6). Decoding performance is significantly above chance in all regions. *p < 0.05 in Wilcoxon signed-rank test. **d,** Trial-by-trial goal-pattern decoding performance aligned to block transitions. Decoding performance was significantly above the theoretical chance level of 0.5 (*p < 0.05, Wilcoxon signed-rank test). During the first trial after each transition (shaded), water was pre-filled at the goal wells and LEDs beneath the maze were illuminated to facilitate recognition of the new goal configuration. Robust goal-pattern decoding was observed from subsequent trials. **e,** Goal-pattern decoding performance during the running phase of the task for each brain region. Decoding performance remains stably above shuffle level across maze positions. **f,** Goal-pattern decoding performance during the stationary (well-licking) phase of the task. Despite marked reductions in running speed and LFP theta power during this phase, goal-pattern decoding remains stable, indicating behavioral-state-independent goal-state coding across the mPFC–NR–CA1 circuit. **g,** Goal-pattern classification based on pre-navigation CA1 spike sequences during quiescent periods. The same classifier as in **a** was applied to spike-sequence events detected in Fig. 2a. Bottom left: Goal-pattern classification for individual spike-sequence events. Right: Box plots showing decoding performance across all 25 sessions (p < 0.001) and individual rats (p = 0.004). Wilcoxon signed-rank test.

Despite this state-agnostic training procedure, goal patterns were reliably decoded throughout the session in all three regions (Fig. 4b). Decoding performance was significantly above chance in mPFC, NR, and CA1 (mPFC: 0.834 ± 0.011; NR: 0.828 ± 0.017; CA1: 0.803 ± 0.013; p < 0.001, Wilcoxon signed-rank test; mPFC: 13 sessions from 3 animals; NR: 12 sessions from 3 animals; CA1: 25 sessions from 9 animals; Extended Data Fig. 7). At block transitions, when the middle-goal location changed, goal-state representations switched within 1–2 trials (corresponding to 4–8 journey segments) across all three regions (Figs. 4d and Extended Data Fig. 7), indicating rapid and coordinated updating of goal-state representations throughout the circuit.

We next examined whether goal-state decoding performance depended on the animal’s behavioral state. Transitions between maze running and stationary well-licking periods were accompanied by pronounced reductions in running speed and hippocampal theta power. Despite these changes, goal-state decoding performance remained stable across behavioral states in all three regions (Fig. 4f). This result indicates that goal-state representations in the mPFC–NR–CA1 circuit are maintained irrespective of locomotor state.

Given the consistent goal-state representations observed during stationary periods, we asked whether these state differences are directly reflected in the brief spike sequences observed in hippocampal CA1 during immobility. To test this, we computed population firing rates during individual pre-navigation spike-sequence events and applied the LDA classifier trained on 2-s segments (Fig. 4g; see Methods). Spike-sequence events were classified according to the animal’s currently engaged goal pattern significantly better than chance (decoding performance: data, 0.604 ± 0.007; chance, 0.553 ± 0.007; p < 0.001, Wilcoxon signed-rank test).

Together, these results demonstrate that goal-state information is persistently represented across mPFC, NR, and CA1 throughout task engagement, independent of whether the animal is locomoting or immobile. This persistent goal-state signal is also expressed in CA1 pre-navigation spike sequences, providing a link between global contextual representations and the transient internal dynamics that precede goal-directed navigation.

### NR silencing impairs goal-dependent CA1 dynamics while sparing spatial coding

The observation that goal states are consistently represented across mPFC, NR, and CA1 suggests that goal-related information is transmitted to CA1 via the mPFC–NR pathway. To directly test this hypothesis, we performed optogenetic silencing of NR neurons while recording CA1 activity during task performance. An adeno-associated virus encoding the inhibitory opsin SwiChR++^50^ was injected into NR, and an optic fibre was implanted at the same site (Figs. 5a and Extended Data Fig. 2). The same animals were also implanted with tetrode microdrives targeting dorsal CA1. NR inactivation was achieved by delivering brief 473-nm laser pulses at low frequency (0.2 Hz) throughout the recording session.

**Figure 5:**
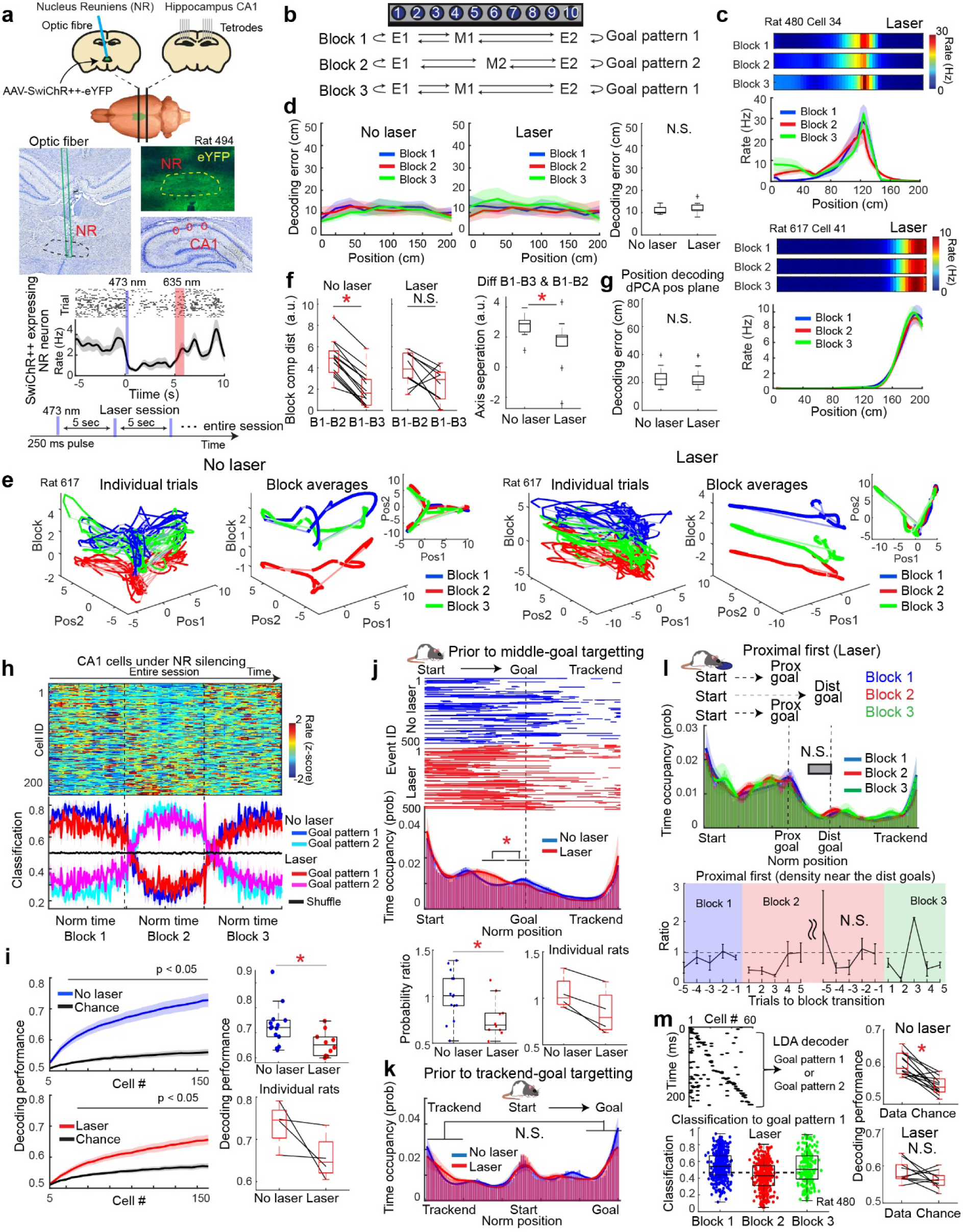
NR silencing diminishes CA1 goal-state coding and pre-navigation prospective activity while preserving spatial representations. **a,** Top: Schematic of the surgical procedures. Middle: Brain sections showing the optic fiber track (top left) and SwiChR++–eYFP expression (top right) in NR, as well as tetrode tracks in CA1 (bottom right). Bottom: Example NR neuron expressing SwiChR++ showing reduced firing in response to 473-nm light delivery. This experiment was performed in a separate animal implanted with an optic fiber attached to a Neuropixels probe in NR. The bottom schematic illustrates the laser application protocol, in which 250-ms pulses of 473-nm light were delivered intermittently at 5-s intervals. **b,** Schematic of the task structure. **c,** Firing-rate maps of two representative CA1 place cells, showing stable spatial tuning with minimal firing-rate modulation across blocks. **d,** Decoding errors across maze positions from CA1 population activity with and without NR silencing. The box plot on the right compares decoding performance between laser and no-laser sessions, showing no significant difference. p > 0.05, Wilcoxon rank-sum test. **e,** CA1 population activity projected onto dPCA-derived orthogonalized block and position axes, as in Fig. 1j, shown for the same animal (rat 617) with and without laser application. While trajectories along the position axes (top right insets) remain consistent across blocks in both conditions, separation of block 2 along the block axis is markedly reduced during NR silencing. **f,** Quantification of block-axis separation between blocks 1 and 2 relative to blocks 1 and 3. The block-specific separation observed in control sessions is largely abolished during NR silencing. No laser: p < 0.001; laser: p = 0.065, Wilcoxon signed-rank test. Right: direct comparison of block-axis separation with and without laser application reveals a significant reduction during NR silencing. p = 0.013, Wilcoxon signed-rank test. **g,** Spatial information occupied in the dPCA-derived position components, assessed using decoding analysis. No significant difference is observed between laser and no-laser sessions. p > 0.05, Wilcoxon signed-rank test. **h,** Top: Color-coded, z-scored activity of 200 randomly sampled CA1 neurons during NR silencing across blocks. Bottom: Comparison of goal-pattern decoding performance from 150 randomly sampled CA1 neurons between laser and no-laser sessions. **i,** Cell-number–matched comparison of goal-pattern decoding performance between laser and no-laser sessions. Performance is compared to corresponding chance levels (Extended Data Fig. 10). Goal-pattern decoding is significantly reduced during NR silencing. p = 0.015, Wilcoxon rank-sum test. **j,** Comparison of time-occupancy probabilities of pre-navigation spike sequences during quiescent periods preceding middle-goal–directed trials, with and without NR silencing. The ratio of occupancy in the fourth versus third quadrants of the goal-directed trajectory is significantly reduced during laser sessions. Bottom box plots summarize data from individual sessions (no laser, n = 14; laser, n = 10) and from 4 rats. *p = 0.021, Wilcoxon signed-rank test. **k,** Same analysis as in **j**, but for trials targeting trackend goals. No significant difference is observed in the ratio of occupancy near the upcoming goal versus the previous goal. p > 0.05, Wilcoxon signed-rank test. **l,** Same analysis as in Fig. 2f, applied to sessions with NR silencing. Time-occupancy probabilities of pre-navigation spike sequences are shown for trials in which animals targeted a proximal goal in blocks 1 and 3 and a distal goal in block 2. Bottom panel shows trial-wise densities relative to block 1. No significant differences are observed across blocks. p > 0.05, Wilcoxon signed-rank test. **m,** Goal-pattern classification based on pre-navigation spike sequences during quiescent periods, as in Fig. 4g. Bottom left: Classification of individual spike-sequence events from the same animal as shown in Fig. 4g (rat 480) during laser sessions. Right: Box plots summarizing decoding performance across no-laser (n = 14) and laser (n = 10) sessions demonstrate that goal-pattern classification is significantly impaired during NR silencing. No laser: p < 0.001, laser: p = 0.193, Wilcoxon signed-rank test.

We first assessed whether NR silencing affected behavioral performance. Comparing sessions with and without laser stimulation, NR inactivation did not significantly alter task accuracy (no laser: 86.5 ± 2.1%; laser: 86.5 ± 3.0%; p = 0.519, Wilcoxon rank-sum test) or running speed (no laser: 20.7 ± 1.1 cm/s; laser: 19.2 ± 1.1 cm/s; p = 0.429; Extended Data Fig. 8). These results indicate that NR silencing did not impair overall task engagement or locomotion.

We next examined the effects of NR silencing on CA1 neuronal activity. We compared recordings from the same animals in sessions with and without laser stimulation (no laser: 923 neurons; laser: 916 neurons; 4 animals). Classification of CA1 neurons revealed a significant reduction in the proportion of rate-remapping place cells during laser sessions compared with control sessions (no laser: 39.1 ± 0.3%; laser: 29.6 ± 2.7%; p = 0.020, Friedman test; Extended Data Fig. 9). A similar reduction was observed for non-spatial cells exhibiting significant goal-dependent rate changes (no laser: 30.3 ± 1.4%; laser: 21.3 ± 1.8%; p = 0.020). In contrast, the proportion of global-remapping place cells did not differ significantly between conditions. Consistent with these findings, population-vector (PV) correlation analyses showed that, under NR silencing, ensemble activity no longer differed significantly between blocks with different versus identical goal configurations for either spatial or non-spatial cells (Extended Data Fig. 9). This contrasted with the clear goal-dependent differences observed in control sessions (Extended Data Fig. 9).

Importantly, NR silencing did not affect the spatial information content of CA1 neurons (no laser: 0.580 ± 0.014 bits/spike; laser: 0.578 ± 0.013 bits/spike; p = 0.934, Kolmogorov–Smirnov test; Extended Data Fig. 9) or position decoding accuracy from CA1 population activity (no laser: 10.9 ± 0.5 cm; laser: 11.9 ± 0.8 cm; p = 0.169; Wilcoxon rank-sum test; Figs. 5d and Extended Data Fig. 9).

To assess how NR silencing altered the structure of CA1 population activity, we applied the same dPCA-based manifold analysis used in Fig. 1. Under NR silencing, the separation between CA1 population trajectories associated with different goal blocks was significantly reduced along the block axis (Figs. 5e–g). This reduction was consistent across sessions and animals (goal-axis separation difference between blocks 1–2 versus blocks 1–3: no laser, 2.81 ± 0.18; laser, 1.64 ± 0.57; p = 0.0128, Wilcoxon rank-sum test). In contrast, spatial representations along the position axes were unaffected, as indicated by unchanged position decoding errors (position decoding error: no laser, 23.5 ± 1.9 cm; laser, 22.9 ± 2.3 cm; p = 0.567).

We further quantified the impact of NR silencing on goal-state information by constructing LDA-based goal-pattern classifiers from randomly selected 2-s population activity segments, irrespective of behavioral state (Fig. 5h). After matching population size across conditions by subsampling, NR silencing significantly reduced goal-pattern decoding accuracy in CA1 (no laser: 72.8 ± 2.2%; laser: 65.6 ± 1.6% with 150 randomly sampled neurons; p = 0.016; Figs. 5i and Extended Data Fig. 10).

Finally, we examined whether NR silencing affected pre-navigation CA1 spike sequences, which normally exhibit goal-dependent modulation (Fig. 2). NR inactivation did not significantly alter the frequency or duration of detected spike-sequence events (no laser: 3.71 ± 0.34 Hz, 51.8 ± 2.0 ms; laser: 3.58 ± 0.34 Hz, 55.1 ± 2.9 ms; p > 0.05 in Wilcoxon rank sum test; Extended Data Fig. 8). However, prior to middle-goal–directed journeys, NR silencing significantly reduced the density of pre-navigation sequences near the goal location (Fig. 5j). In contrast, the spatial distribution of pre-navigation sequences preceding trackend–goal–directed journeys was largely unaffected (Fig. 5k). Moreover, under NR silencing, pre-navigation sequence activity no longer adapted to changes in middle-goal location across blocks (Figs. 5l and Extended Data Fig. 10), indicating that the impact of NR silencing on goal-directed sequences is specific to target wells that change between blocks.

Consistent with these observations, classification of individual pre-navigation spike-sequence events by goal pattern dropped to chance levels during NR silencing, whereas significant classification performance was observed in control sessions (decoding performance: no laser, 0.634 ± 0.019; laser, 0.583 ± 0.006; chance, 0.568 ± 0.009; p = 0.011 no laser vs. chance, p =0.131 laser vs. chance in Wilcoxon signed rank test; Fig. 5m). Together, these results demonstrate that NR input is required for transmitting goal-state information to CA1, enabling goal-dependent modulation of CA1 population activity and pre-navigation internal dynamics while leaving spatial coding largely intact.

## Discussion

In this study, we show that prefrontal top-down input routed through the mPFC–NR–CA1 pathway enables the hippocampus to distinguish distinct navigation episodes within the same environment while preserving a stable spatial map. CA1 achieves this through an orthogonal coding scheme in which spatial position and goal-state information occupy largely independent dimensions of the population activity manifold. This organization allows goal-related signals to be incorporated without disrupting positional coding, thereby minimizing representational interference between episodes formed in the same spatial context. Importantly, goal-state signals persist across both locomotor and stationary phases, enabling the hippocampus to engage an episode-specific map not only during navigation but also during planning before movement begins.

Previous studies have implicated the prefrontal cortex in representing navigational goals or intended destinations^36,40^, raising the possibility that prefrontal–hippocampal interactions shape goal-dependent hippocampal activity. Here, however, we find that mPFC and NR do not encode the immediate next goal of individual journeys (Extended Data Fig. 6). Instead, these regions form persistent neural states that reflect the block-wise goal configuration. We thus use goal state to refer to the block-wise configuration of rewarded locations, not the next destination within a journey. These states are maintained across locomotion and immobility and remain robust despite trial-by-trial behavioral variability, suggesting that they provide a stable contextual signal that distinguishes goal-configuration–specific episodes over extended timescales. This contextual signal is relayed via NR to CA1, where it selectively modulates the goal-coding dimension of the CA1 population manifold. Consistent with this model, inactivation of NR disrupted CA1 goal-state coding, reduced separation between episode-specific CA1 manifolds, and impaired goal-dependent prospective activity in pre-navigation CA1 spike sequences, indicating that this prefrontal–thalamic input is required for robust expression of the goal-state dimension. This selective dissociation, disrupting goal-state coding while sparing spatial representations, provides causal evidence that prefrontal–thalamic input does not reshape the hippocampal spatial representations, but instead controls which episode-specific map is engaged.

These findings extend classical views of rate remapping in the hippocampus, in which place cells alter their firing rates across contexts while preserving their spatial tuning. Classical rate remapping is typically driven by changes in sensory input and is associated with active locomotion^10^. In contrast, the goal-state representations described here reflect internal contextual and planning-related signals rather than externally perceived sensory or spatial information, and they persist across both running and quiescent behavioral states. This behavioral-state independence is notable because it contrasts with the strong locomotion dependence observed in most hippocampal activity patterns, including large changes in firing rates and local field potential theta power^32–35^. By comparison, goal-state coding in mPFC and NR during stationary phases is less sensitive to behavioral-state-dependent activity (Extended Data Fig. 7), potentially enabling these regions to provide CA1 with a stable goal-state signal across behavioral states.

This persistence of goal-state representations offers a mechanistic explanation for how the hippocampus can generate goal-dependent pre-navigation spike sequences. Because goal-specific CA1 population states are already engaged during immobility, the hippocampus can express prospective activity biased toward the intended destination before movement onset. This framework helps reconcile diverse observations regarding hippocampal pre-navigation spike sequences, which have been reported to represent future, past, or alternative trajectories depending on task demands^26,27,51^. Such variability may reflect top-down control over which episode-specific map is selected at a given moment, biasing the generation of behaviorally relevant sequences, as proposed in several theoretical models^52,53^.

More broadly, this mechanism offers a circuit-level solution to how memory systems achieve flexible retrieval and planning in stable environments^2,54–57^ —a problem central to episodic memory, cognitive control, and adaptive behavior across species. Our findings suggest that the hippocampus resolves this challenge by constructing parallel spatially aligned maps, each corresponding to a distinct behavioral episode through orthogonal coding of spatial and non-spatial information. A key computational consequence of this architecture is that previously established maps can be preserved rather than overwritten, as new experiences accumulate. When a previously encountered goal pattern reappeared after an intervening block, the corresponding CA1 population representation was selectively reinstated, indicating that CA1 maintains multiple episode-specific maps within the same environment. This property is a defining feature of a memory system and is distinct from the gradual temporal drift of neural representations^58–60^. Although CA1 activity exhibited temporal drift, this drift occurred along a dimension largely independent of the goal-state axis, preserving episode-specific separability (Extended Data Fig. 10). Such an organization allows the hippocampus to retain stable representations of shared structure while flexibly indexing individual episodes. In this framework, prefrontal contextual input transmitted via the nucleus reuniens can be viewed as a selection signal that biases which episode-specific map is engaged at a given time. This interpretation is consistent with the proposed roles of the prefrontal cortex^61–63^ and nucleus reuniens^64,65^ in memory retrieval and consolidation. Notably, this prefrontal–thalamic signal persists across both locomotion and immobility, enabling the hippocampus to maintain a consistent internal state that links task-specific activity across movement and stationary phases, including rate-remapping and replay. By identifying a prefrontal–thalamic mechanism that selects among spatially aligned hippocampal maps, our findings bridge circuit-level interactions and population-coding principles that support flexible memory access, planning, and goal-directed behavior.

## Materials and Methods

### Subjects

All experiments were approved by the local authorities (RP Darmstadt, protocols F126/1009, F126/1026, and F126/2005) in accordance with the European Convention for the Protection of Vertebrate Animals used for Experimental and Other Scientific Purposes. Subjects were fifteen male Long-Evans rats weighing 400 to 550 g (aged 3–5 months) at the start of the experiment. Rats were housed individually in Plexiglass cages (45 × 35 × 40 cm; Tecniplast GR1800) and maintained on a reversed 12 hr light-dark cycle, with behavioral experiments performed during the dark phase. Animals were mildly water-restricted with unlimited access to food and kept at 85–90% of their free-feeding body weight throughout the experiment. Nine rats were implanted with tetrode drives bilaterally in the dorsal CA1 of the hippocampus. Four of them additionally received injections of AAV2/1-CamKIIα-SwiChR++-eYFP into the nucleus reuniens (NR) of the thalamus, where optic fibers were also implanted. Five rats were implanted with Neuropixels 1.0 probes: one with 2 probes in NR, one with 1 probe in NR, two animals with 1 probe in mPFC, and another with 1 probe in mPFC and 2 probes in NR.

### Surgery, virus injection, and drive implantation

Anesthesia was induced by isoflurane (5% induction concentration, 0.5–2% maintenance adjusted according to physiological monitoring). For analgesia, Buprenovet (Buprenorphine, 0.06 mg/mL; WdT) was administered by subcutaneous injection, followed by local intracutaneous application of either Bupivacaine (Bupivacaine hydrochloride, 0.5 mg/mL; Jenapharm) or Ropivacaine (Ropivacaine hydrochloride, 2 mg/mL; Fresenius Kabi) into the scalp. Rats were subsequently placed in a Kopf stereotaxic frame, and an incision was made in the scalp to expose the skull. After horizontal alignment, several holes were drilled into the skull to place anchor screws, and craniotomies were made for microdrive implantation. The microdrive was fixed to the anchor screws with dental cement, while two screws above the cerebellum were connected to the ground of the recording devices. All tetrodes were then positioned at a depth of 750 μm from the cortical surface. All animals received analgesics (Metacam, 2 mg/mL Meloxicam; Boehringer Ingelheim) and antibiotics (Baytril, 25 mg/mL Enrofloxacin; Bayer) for at least 5 d post-surgery.

For tetrode recordings, rats were implanted with a microdrive that contained 28 individually adjustable tetrodes made from 17 μm polyimide-coated platinum-iridium (90–10%; California Fine Wire; plated with gold to impedances below 150 kΩ at 1 kHz). The tetrode bundle consisted of 30-gauge stainless steel cannulas, soldered together in 14 × 2 circular shapes. Tetrode drives were implanted within the dorsal CA1 region of the hippocampus bilaterally, with coordinates at AP: −4.0 mm, ML: 3.5 mm. The tetrodes were adjusted to depths of 1.8– 2.2 mm DV. Concurrently, four of these rats also received injections of AAV2/1-CamKIIα-SwiChR++-eYFP (a gift from Dr Karl Deisseroth)^50^ into NR at four sites in each hemisphere, with each site receiving 250 nl (a total of 1000 nl per hemisphere). The injections were administered at a rate of 100 nl/min with coordinates of AP: 2.0 mm and 2.5 mm, ML 0.6 mm, and DV 6.5 and 6.75 with an angle of 4° toward the midline. After each injection, there was a 10-minute waiting period before proceeding with the next injection site. An optic fiber (lambda fibre, OptogeniX) was implanted into the NR at the coordinates of AP: 2.25 mm, ML 0.6 mm, and DV: 6.75 mm with an angle of 4° toward the midline. After the surgery, the animals were allowed to recover for 5-7 days before starting recordings. Optogenetic experiments were conducted at least 4 weeks after surgery to allow for opsin expression.

In addition, one rat was implanted with a Neuropixels 1.0 probe in NR with coordinates of AP 2.5 mm, ML 0.6 mm, and DV: 7.5 with an angle of 5° to the midline. Another subject was implanted with two Neuropixels probes in NR bilaterally with the aforementioned coordinates. One rat had three Neuropixels 1.0 probes: two in NR bilaterally and another in the mPFC with the coordinates of AP 3.25 mm, ML 0.6 mm, and DV 5.0 mm with an angle of 4 degrees to the midline. Two rats were implanted with one Neuropixels 1.0 probe in the mPFC.

### Histological procedures

Once the experiments were completed, the animals were deeply anesthetized with sodium pentobarbital and perfused intracardially with saline, followed by 10% formalin. The brains were extracted and fixed in formalin for at least 72 h at 4 °C. Frozen coronal sections were cut (30 µm), stained with crystal violet, and mounted on glass slides. To confirm the implanted locations of the tetrodes, optic fibers, or Neuropixels probes, coronal sections at the implanted locations were directly mounted on glass slides from the cryostat, stained with cresyl violet, and covered with coverslips using mounting solution (HISTOMOUNT, Invitrogen, 008030).

To confirm AAV-mediated protein expression, sections were immunostained to enhance the fluorescence signals of SwiChR++-eYFP. The sections were first placed in a blocking buffer (4% [v/v] normal goat serum and 2% [w/v] bovine serum albumin in pH 7.6 TBST) for 30 min. The staining of eYFP was performed with an antibody against GFP (anti-GFP Chicken IgY, 1:1000 dilution, Invitrogen, A10262) together with a corresponding secondary antibody (anti-Chicken Goat IgY AF594, 1:500 dilution, Invitrogen, A32759). All the images were taken using a slide scanner (Zeiss Axio Scan.Z1).

### Behavioral task

Rats were acclimated to a 2-m-long linear maze equipped with ten equally spaced water-delivery wells (20 cm apart). The training regimen was composed of three phases. Initially, rats were familiarized with the maze by having access to 100 µl liquid rewards (0.3% saccharin) positioned at both ends of the maze. Rewards were prefilled in the wells, facilitating the animals’ learning to alternate between the trackend wells over the first one to two days. In the second phase, a middle well was introduced to the goal-well sequence. Rewards were prefilled in the designated wells to facilitate the animal’s learning of a three-well navigation sequence. This training phase took 1–2 days. Finally, a time delay between an animal’s well licking and the reward delivery was introduced. The duration of this delay was fixed throughout each behavioral session, but gradually increased over a period of 7 to 14 days until it reached a range of 0.8 to 1.5 seconds. Once rats performed the task with a minimum delay of 0.5 seconds over 80% accuracy, the middle well in the sequence was altered to a new location after several trial repetitions. To signal this transition, a reward was prefilled at a new goal well and LEDs underneath all wells in the maze were turned on until the completion of the first cycle of new goal sequence trials (comprising four journey segments). All the experiments were performed under minimum light conditions (no light source in the recording room, with only weak ambient light from the computer monitors in the adjacent room), and the animal’s position and direction were monitored by two-colored LEDs on the headstage.

### Optogenetic Inactivation

Optogenetic experimentation commenced after three weeks of these recording sessions (approximately four weeks post-surgery). During these experiments, 473 nm laser pulses of 250 ms duration were emitted at a frequency of 0.2 Hz using a DPSS laser (Shanghai Laser & Optics Century Co., Ltd.). The laser power at the fiber tip was adjusted to 20 mW. These pulses were applied throughout the recording session. After the session, 635 nm light from a diode laser (Shanghai Laser & Optics Century Co., Ltd) was applied for 10 s to facilitate the recovery from SwiChR++-dependent inactivation.

### Recording procedures and spike sorting

All data processing and analyses were performed with MATLAB (MathWorks). Neural signals from tetrode recordings were acquired and amplified using 64-channel headstages (RHD2164; Intan Technologies), combined with the OpenEphys acquisition system at a sampling rate of 15 kHz. The signals were band-pass filtered at 300–6000 Hz, and spikes were detected and assigned to separate clusters using Kilosort^66^ (https://github.com/cortex-lab/KiloSort) with parameters set to the spike threshold at −4 and the number of filters at 2× the total channel number. Each channel was independently grouped with ‘kcoords’ parameters, and the noise parameter determining the fraction of noise templates spanning across all channel groups was set to 0.01. For Neuropixels probe recordings, the signals, high-pass filtered at 300 Hz, were collected with the National Instruments PXIe System and OpenEphysGUI software. Spikes were detected and assigned to separate clusters using Kilosort 3.0 (https://github.com/MouseLand/Kilosort). The obtained clusters were checked and adjusted manually based on autocorrelograms and waveform characteristics in principal component space, obtaining well-isolated single units by discarding multi-unit activity or noise. Neurons with mean firing rates less than 0.1 Hz were excluded.

### Cell classification

To assess the spatial tuning properties of CA1 neurons during navigation, we first defined the beginning of individual journeys using the criteria of the animal’s running speed > 10 cm/s and position away from the start well > 10 cm. The end of the journey was defined as the time of the first lick detection at the goal well. The error and transition trials (the first cycle with new goal introduction) were excluded. We considered firing rates on these defined navigation phases from the beginning to the end of journeys. To avoid multiple counts of the same cells, we used only one recording session from each animal. Spike times of individual CA1 neurons were converted into firing rates by convolving a Gaussian kernel with a bandwidth of 250 ms. We then computed the spatial information content of each neuron’s firing pattern. The spatial information metric quantifies how well spikes of a neuron predict the animal’s location in the environment. We adopted the formula provided by Skaggs et al. ^67^, which accounts for the variability in firing rate across different spatial bins and the occupancy probability of each bin, as shown in the following formula:

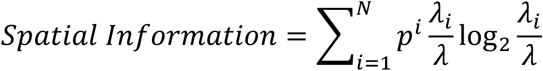

Here, *pi* is the occupancy probability of the ith bin, λ_*i*_ is the mean firing rate at the i-th bin, λ is the overall mean firing rate, and *N* is the number of bins. All neurons that exhibited the maximum firing rates above 1 Hz across the two blocks with the same goal combinations (i.e., blocks 1 and 3 in Fig. 1a) were considered. Among them, neurons that exhibited spatial information > 0.25 bits/spike in both blocks were considered spatial cells, and the rest were considered nonspatial cells.

Spatial cells were further categorized based on the stability of spatial tuning. The significance of spatial tuning differences was assessed based on spatial correlations between 1-dimensional firing rate maps along the linear maze obtained from two blocks for each neuron. By shifting the relative positions of these rate maps along the maze positions, we obtained the distributions of spatial correlations across different spatial alignments of the two rate maps. If the 95th percentile of this distribution can be achieved within a 10 cm shift of the original data, these two maps were considered to have stable spatial tuning between blocks. Spatial cells with stable spatial tuning between two same-goal blocks were considered place cells.

Place cells were further classified based on their remapping features. Global-remapping cells were defined as neurons that exhibited significant field shifts between blocks with different goal combinations (i.e., blocks 1 and 2) by using the same spatial correlation criteria described in the previous paragraph. For the remaining place cells excluding global-remapping cells, we assessed the overall rate changes by using two-way ANOVA with goal patterns and spatial positions as independent variables. Neurons that showed p < 0.05 in the main goal-pattern effect of ANOVA and exhibited larger rate differences between the different goal blocks (blocks 1 and 2) than the same ones (blocks 1 and 3) were considered rate-remapping place cells.

The same criteria for rate remapping were applied for nonspatial CA1 cells. Nonspatial neurons that exhibited p < 0.05 in the main goal-difference effect of ANOVA and exhibited larger mean rate differences between the different goal blocks than the same ones were classified as nonspatial rate-remapping cells.

The centre of mass (COM) of firing fields in the maze was calculated for individual neurons using the following formula:

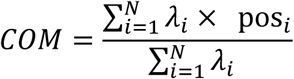

Here, λ_*i*_ is the mean firing rate at the i-th bin, pos_i_ is the spatial position of the i-th bin, and *N* is the number of bins. Changes in spatial tuning were assessed by computing the absolute differences in the COMs between blocks. The rate changes between blocks were defined as the absolute difference between the peak firing rates in the two blocks, normalized by their maximum. The comparison of the distributions was performed using the Kolmogorov-Smirnov test.

### Population vector and statistical analysis

Previous studies have shown that, even when individual neurons maintain the same spatial tuning, spatial codes at a neural population level can differ by changing the distribution of firing rates among neurons (i.e., rate remapping)^10^. To evaluate this effect, we performed a population vector (PV) correlation analysis on all place cells, excluding global-remapping cells. The z-scored firing rates of neurons were calculated for individual 2 cm position bins. Correlations of population firing rates across position bins between different blocks were calculated. Statistical analyses were performed on the PV correlations along the same positions. We used only one recording session per animal, but datasets from two different running directions were treated as repeated measures from the same animals.

### Position Decoding

To estimate the spatial position represented by CA1 neurons, we applied a Bayesian decoding approach with a uniform prior^26,45^. This approach requires constructing spatial rate maps for individual neurons. Spike times of individual CA1 neurons were converted into firing rates by convolving a Gaussian kernel with a bandwidth of 250 ms, and the mean firing rates in individual 2 cm bins were estimated across the times when the animal occupied these bins. The times when the animal’s running speed was < 10 cm/s were excluded. We constructed separate maps for the two running directions in the linear maze. By using these rate maps, position decoding was performed from the population activity of CA1 neurons. We assumed that neurons exhibit Poisson firing statistics, and then, the likelihood of observing a spike rate vector at individual positions and directions is given by:

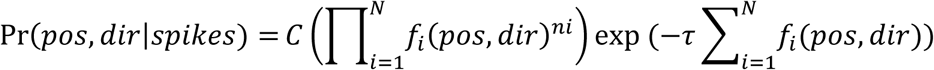

where *C* is a normalization constant, τ is the time bin width, *f*_*i*_ (*pos, dir*) is the mean firing rate of a neuron i at a given position and direction, *n*_*i*_ is the observed spike count of the neuron i, and *N* is the total number of neurons. A 250 ms time window was used to estimate the animal’s position on a behavioral timescale. The maximum likelihood of position and direction was estimated from the neural population activity in individual time bins.

To avoid the overfitting bias of a decoder, we assessed the decoder’s performance using a ten-fold cross-validation approach. Briefly, we extracted 10% of the data as a test dataset, and the remaining datasets were used for training to construct rate maps. We repeated the estimations for all the datasets to be tested. The decoding performance was then estimated by calculating the position decoding errors and the coefficient of determination (R^2^) for both position and direction estimates relative to those obtained from a video camera.

To estimate the impact of goal changes on the position decoding performance, we took two different decoding approaches. In the first approach, the data were combined across blocks to train a decoder, and its performance was estimated on test datasets from individual blocks (general decoder). In the second approach, decoders were constructed individually for each block of data, and their performance was evaluated on test datasets from the same blocks (blockwise decoder). We found that the decoding performance was not significantly different between these two strategies nor between blocks (Extended Data Fig. 3).

### Pre-navigation Spike Sequence Detection

To detect hippocampal replay events, we applied the same Bayesian decoder used for position estimation on a behavioral timescale. However, we restricted position decoding to the animal’s stationary times before navigation, focusing on 3-s periods before leaving start wells with running speed below 10 cm/s. We applied a moving time window of 20 ms, shifting every 5 ms, to estimate positions. The decoded positions were then assessed using the following three criteria for replay sequences. First, consecutive decoded positions should not exhibit a sudden jump of more than 20 cm. Second, the duration of sequences must exceed 20 ms. Finally, the decoded sequences should move at least 25 cm away from the animal’s position. The time occupancies of sequence representations were estimated at every 2.5 cm bin for each recording session and normalized for the distances of the middle or trackend goal locations.

### Dimensionality Reduction and Manifold Analysis

To characterize the structure of neural population dynamics jointly encoding goal patterns and spatial positions, we applied demixed principal component analysis (dPCA)^48^. Neural activity was organized by neuron identity, maze position, trial block, and trial. Maze positions were discretized into 20 spatial bins for each running direction. To minimize the influence of behavioral uncertainty during goal adaptation, neural activity from the first three cycles following each change in goal configuration was excluded from the analysis. The dPCA was then applied to this dataset to extract components associated with spatial positions and task blocks. A regularization parameter of 10^-4^ was used to limit the contribution of small eigenvalues, thereby reducing overfitting and improving the robustness of component estimation. From the resulting components, we selected the first block component and the first two position components for further analysis. Because dPCA components are not guaranteed to be mutually orthogonal, we orthogonalized these three components using a Gram–Schmidt procedure, removing projections of the block component from the position components. Population activity from individual trials was subsequently projected onto the orthogonalized components, yielding a low-dimensional representation that preserved task-relevant variance. These components were visualized in three-dimensional space to illustrate the trajectories of neural population activity across different block and spatial conditions.

### LDA-based Goal-pattern and Position Decoding

To quantify spatial and goal-state information across the mPFC–NR–CA1 circuit without assuming spatial tuning of individual neurons, we applied an LDA-based decoding approach to estimate the animal’s position. Neural data were divided into training and test sets using 10-fold cross-validation. Firing rates were estimated by convolving spike trains with a Gaussian kernel (bandwidth 250 ms) and were z-score normalized across the session.

We constructed LDA decoders from ensemble neural activity corresponding to 20 spatial bins for each running direction along the linear maze, separately for each trial block. Using the withheld test data, we first assessed whether goal patterns could be decoded from population activity. We then constructed an LDA decoder for spatial position, yielding 39 (40 − 1) projection axes, and evaluated decoding performance across the full dimensionality on the test dataset. To generate orthogonal decoders for goal and position, we first reduced the dimensionality of the position decoder by retaining components explaining 80% of the variance. From this position-coding subspace, the component of the goal-coding axis was subtracted, resulting in a position subspace orthogonal to goal coding. All LDA decoders were implemented using the MATLAB fitcdiscr function with a regularization parameter gamma of 0.1, and decoding probabilities were obtained using the predict function on the test dataset.

### Goal-pattern Decoding

To decode goal patterns associated with individual task blocks, we applied linear discriminant analysis (LDA) to population neural activity across the recording session. Neural activity from the entire session, including both running and immobility periods, was included in the analysis. The data were segmented into 2-s windows. To construct the goal-pattern decoder, neural activity from the first three cycles following the introduction of a new goal configuration was excluded to avoid contamination by transitional uncertainty during goal adaptation. An LDA decoder was trained using randomly sampled 2-s segments from blocks sharing the same goal pattern. Specifically, 15 segments were sampled from each of blocks 1 and 3 (goal pattern 1) and 30 segments from block 2 (goal pattern 2), thereby equalizing the total number of training samples across goal patterns. The trained decoder was then applied to the remaining 2-s segments from the entire recording session, including segments spanning goal transitions, to estimate the probability that each segment belonged to goal pattern 1 or goal pattern 2. This procedure was repeated 1,000 times with independent random sampling, and decoding performance at each time segment was averaged across iterations to obtain a representative estimate for the session.

Chance-level performance was assessed using two complementary approaches. First, a shuffled-label control was implemented using the same decoding procedure, except that goal-pattern labels were randomly shuffled during decoder training. This procedure was repeated 1,000 times to generate a distribution of chance-level decoding performance. The second control was to estimate whether these classifications were specific to goal changes. This was performed by artificially assigning three consecutive segments within the same goal block to different classes (class 1 for the first and third segments, class 2 for the second segment) and applying the same decoding procedure to these data. This analysis was repeated for each of the three blocks, and decoding performance was averaged across blocks to estimate chance-level performance.

### Goal-Pattern Classification for Spike Sequence Events

To assess whether pre-navigation CA1 spike sequences generated during immobility discriminate between goal patterns, we applied a goal-pattern LDA decoder directly to the population spiking activity during individual spike-sequence events. Whereas goal-pattern decoding described in the previous section was based on firing rates estimated using a Gaussian kernel with a bandwidth of 250 ms, we reduced the kernel bandwidth to 50 ms to accommodate the brief duration of spike-sequence events. For each sequence event, mean firing rates during the event were computed and fed into the LDA decoder to classify the underlying goal pattern.

### Statistical analysis

Statistical analyses were conducted using MATLAB (2024a) statistical toolbox (MathWorks). All the statistical tests were two-sided and non-parametric unless stated otherwise. The plots and text display mean (line) ± s.e.m. (shaded).

