## Extended Data Figures for "Locomotion-invariant prefrontal–thalamic goal states organize spatially aligned episode-specific hippocampal maps"

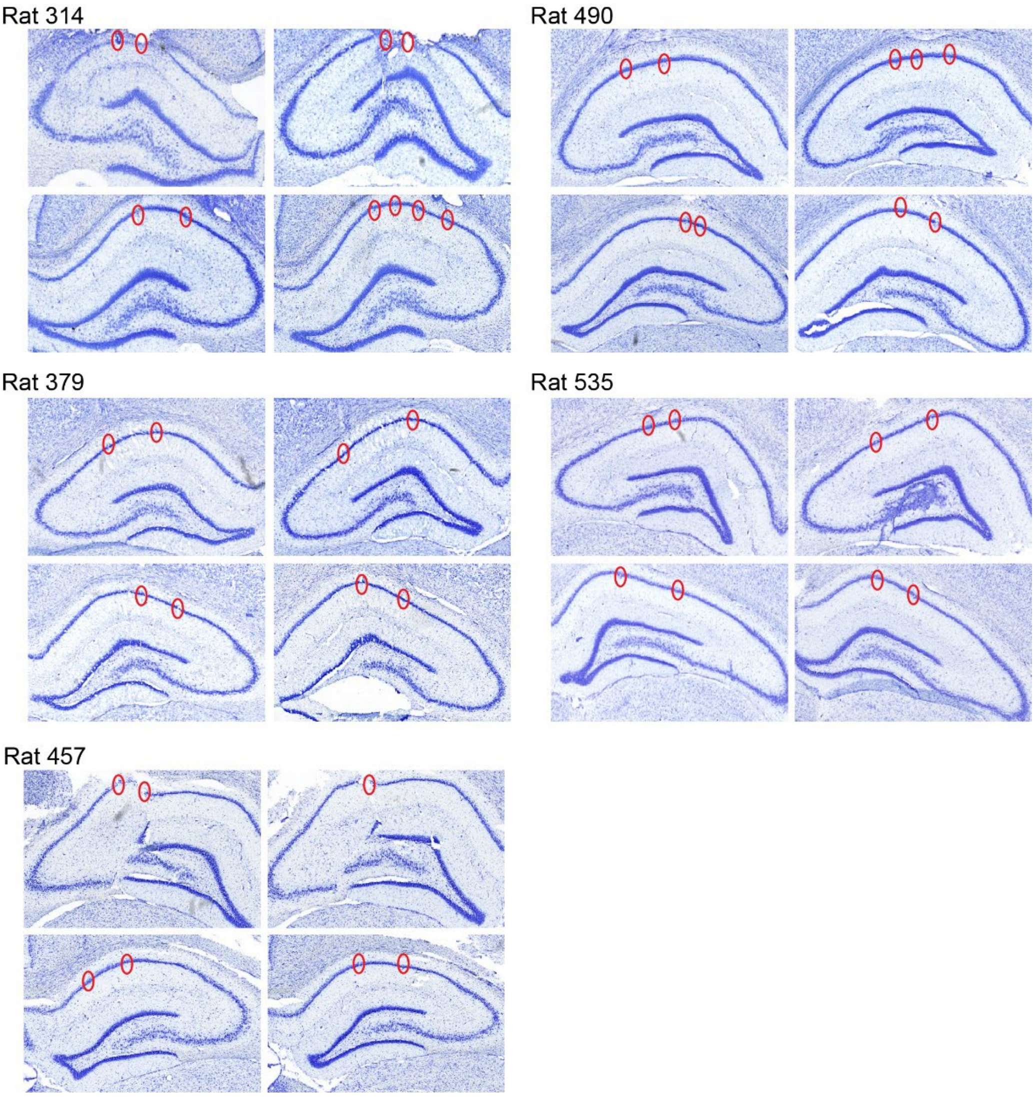


**Extended Data Figure 1: Nissl-stained coronal sections showing electrode positions in the hippocampal CA1.**
Tetrode tracks are marked by red circles. Tetrodes were implanted bilaterally in CA1.

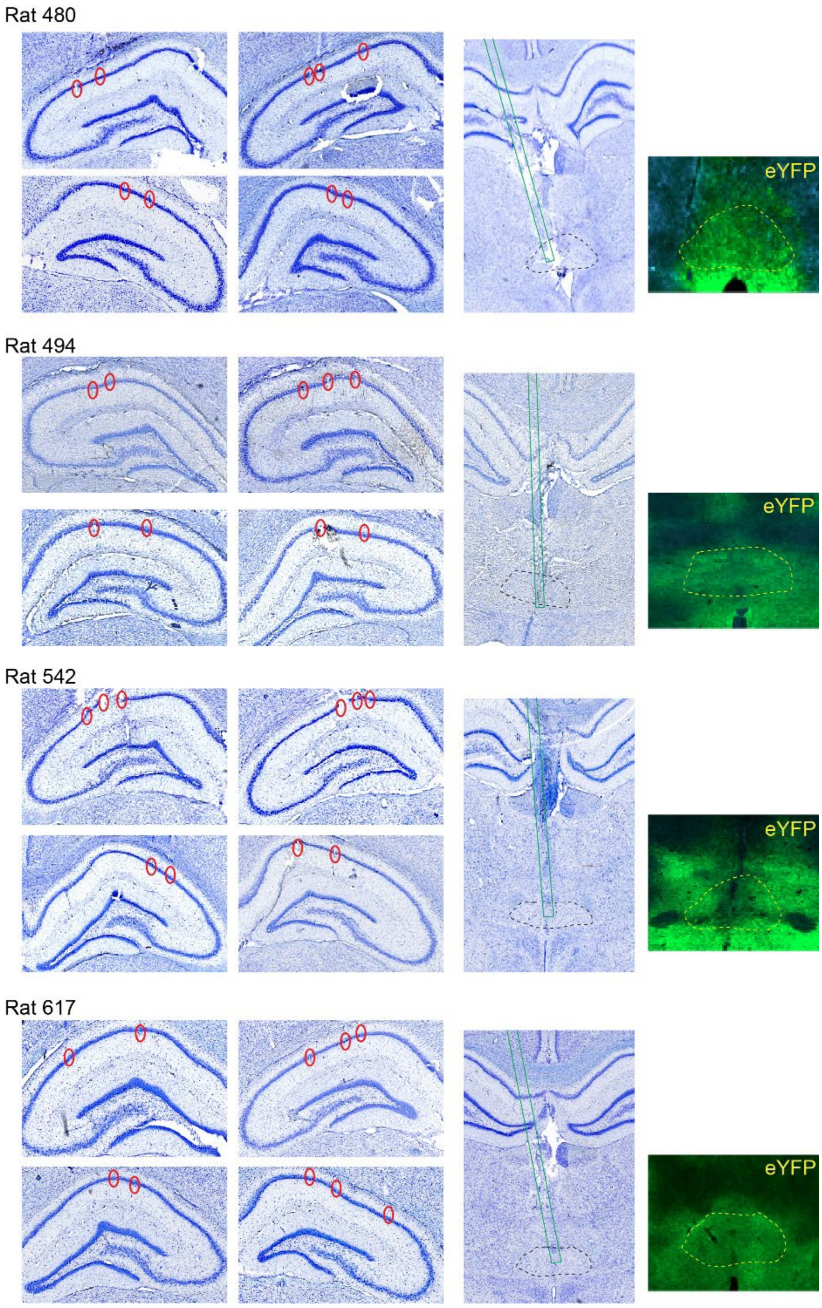


**Extended Data Figure 2: Nissl-stained coronal sections showing electrode locations in the hippocampal CA1, together with optic fiber tracks and SwiChR++ expression in NR.** Brain sections are from animals used for CA1 recordings during optogenetic silencing of NR. Tetrode tracks are indicated by red circles and tetrodes were implanted bilaterally in CA1. For each animal, the two rightmost sections show the optic fiber track (green line) targeting NR (black dashed outline), as well as eYFP immunofluorescence used to verify SwiChR++ expression (green) within NR (yellow dashed outline).

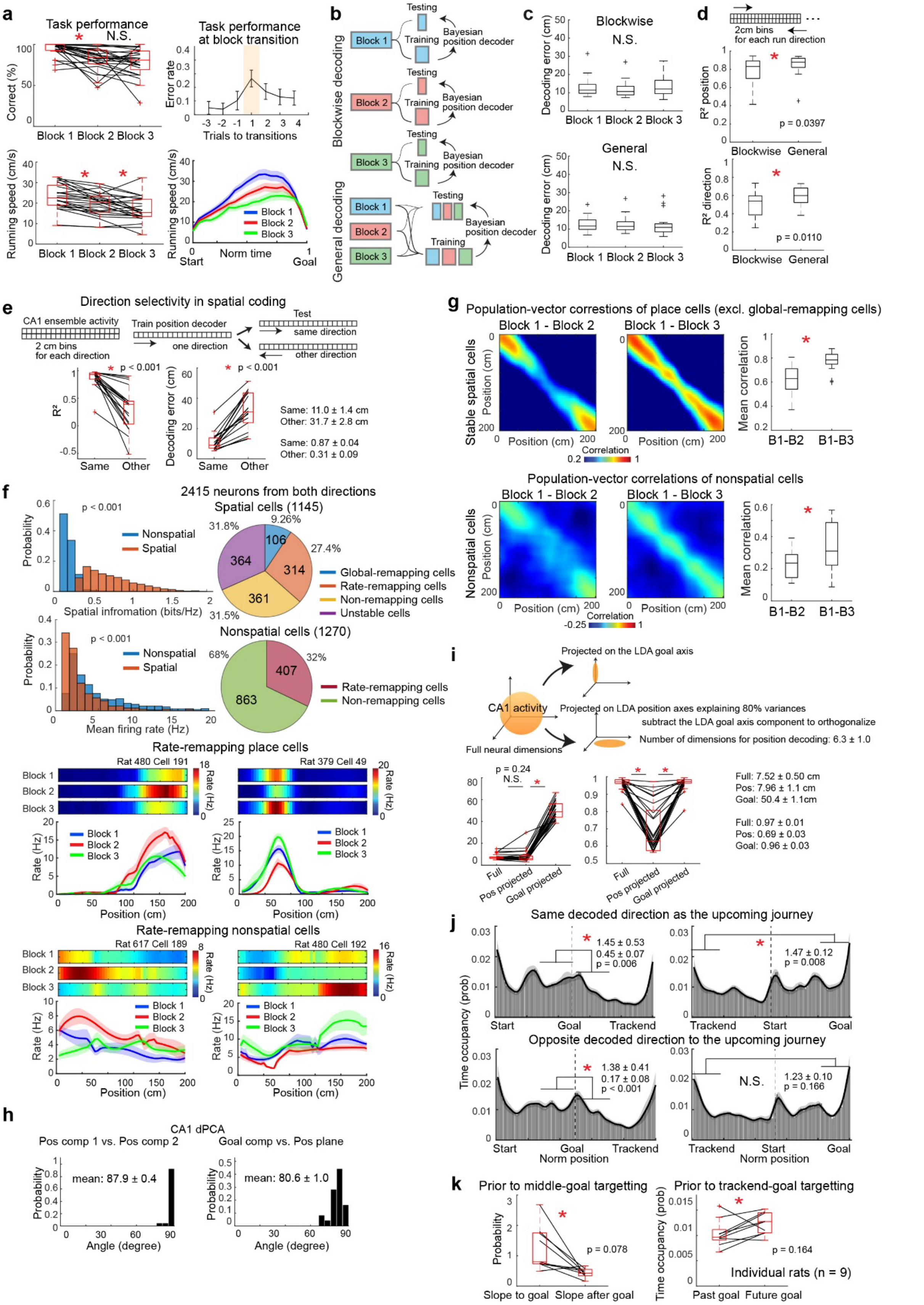


**Extended Data Figure 3: Behavioral analyses, decoding performance, neuronal classification, and pre-navigation CA1 spike sequences.**

**a,** Task performance. The top left panel shows correct performance across blocks, and the top right panel shows error rates at block transitions. Trial 1 denotes the initial journeys introducing a new goal, during which reward water was pre-filled and LEDs beneath all wells were illuminated. The data show that animals rapidly adapt to goal changes. The bottom two panels show running speed for individual journeys from the start to the goal wells, revealing a gradual decrease in running speed over the course of the session. $p < 0.05$, Wilcoxon signed-rank test. **b,** Schematic showing the differences between the blockwise and general decoders. **c,** Left: plots showing position decoding errors across blocks by using the blockwise (top) or general decoder (bottom). Kruskal-Wallis test. **d,** Comparisons of the coefficient of determination ($R^2$) between the two decoders for position and direction estimations. Overall, the general decoder exhibits significantly better performance. 25 sessions from 9 animals. $p < 0.05$, Wilcoxon signed rank test. **e,** Direction dependency of position decoding. The Bayesian position decoder was constructed for neural activity during one running direction, and it was applied to estimate positions when the animal ran in either the same or the other direction. The decoding performance was largely reduced for the opposite direction, suggesting direction-dependency in spatial coding. $p < 0.001$, Wilcoxon signed rank test. **f,** The left two panels show histograms of spatial information and mean firing rates for spatial and nonspatial CA1 neurons. $p < 0.001$, Kolmogorov–Smirnov test. The two pie charts on the right show the proportions of different CA1 cell categories, shown separately for spatial and nonspatial neurons. The bottom panels show representative examples of spatial and nonspatial CA1 neurons exhibiting rate remapping following goal changes. **g,** Population-vector (PV) correlations for place cells excluding global-remapping cells (top) and for nonspatial cells (bottom). Left, color-coded matrices show PV correlations at individual maze positions between blocks 1 and 2, or blocks 1 and 3. Right, box plots show position-averaged PV correlations for each block pair. $p < 0.05$, Friedman test. **h,** Angles between the two position components and between the block component and the two position components extracted by dPCA, demonstrating near-orthogonality of the axes. **i,** Orthogonal representations of position and goal in CA1 population activity. Neural activity was computed in 20 position bins for each of two running directions across three task blocks. A goal classifier was constructed using LDA to discriminate the two goal patterns, yielding a single projection axis corresponding to the goal-coding dimension. An LDA decoder for position was also constructed, yielding 39 ($40 - 1$) projection axes. Dimensionality was reduced by retaining components explaining 80% of the variance ($6.3 \pm 1.0$ dimensions). From this position-coding subspace, the component projected onto the goal-coding axis was subtracted, yielding a position subspace orthogonal to the goal-coding axis. We then compared spatial and goal information using LDA decoding based on the full neural space, the position-coding subspace, or the goal-coding axis alone. Spatial information was largely preserved in the position subspace but not in the goal-coding axis, whereas goal information was confined to the goal-coding axis and largely absent from the position subspace. These results provide evidence that CA1 population activity orthogonalizes position and goal-pattern information within the neural activity manifold. **j,** The Bayesian decoder provides estimates of both position and running direction. We therefore asked whether the direction of decoded positions influences pre-navigation spike sequences. The top two panels show histograms of the temporal occupancy of pre-navigation spatial representations in the same direction as the animal's upcoming journey. The bottom two panels show the corresponding histograms for pre-navigation spatial representations in the direction opposite to the upcoming journey. Overall, the results are similar to those in Fig. 2e. However, a key difference is that prospective coding toward the track-end goals becomes non-significant when only positions in the opposite direction are considered (bottom right). This finding indicates that pre-navigation prospective coding is supported by directional representations aligned with the animal's forthcoming journey. **k,** Supporting analysis for Fig. 2e showing that the results are consistent across all nine recorded animals. $p < 0.05$, Wilcoxon signed-rank test.

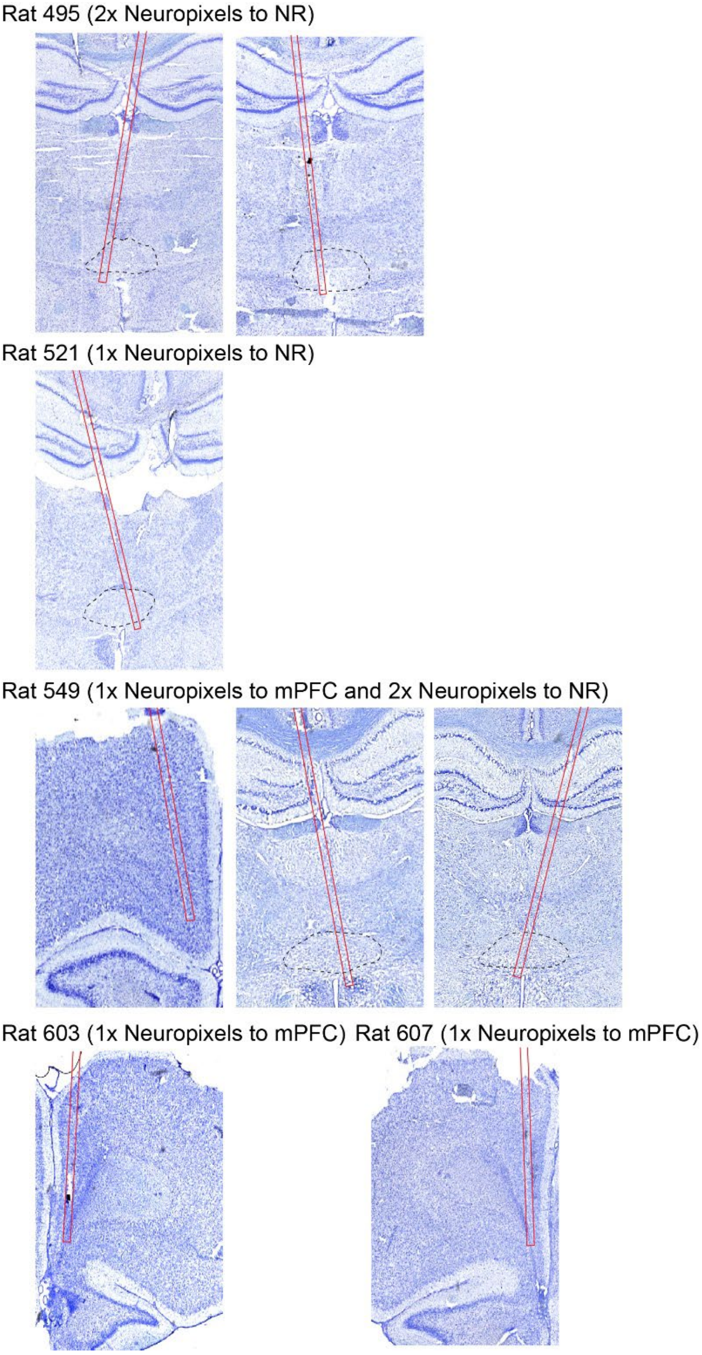


**Extended Data Figure 4: Nissl-stained coronal sections showing the positions of Neuropixels probes in mPFC or NR.**
Probe tracks are indicated by red lines, and the NR is outlined by a black dashed line.

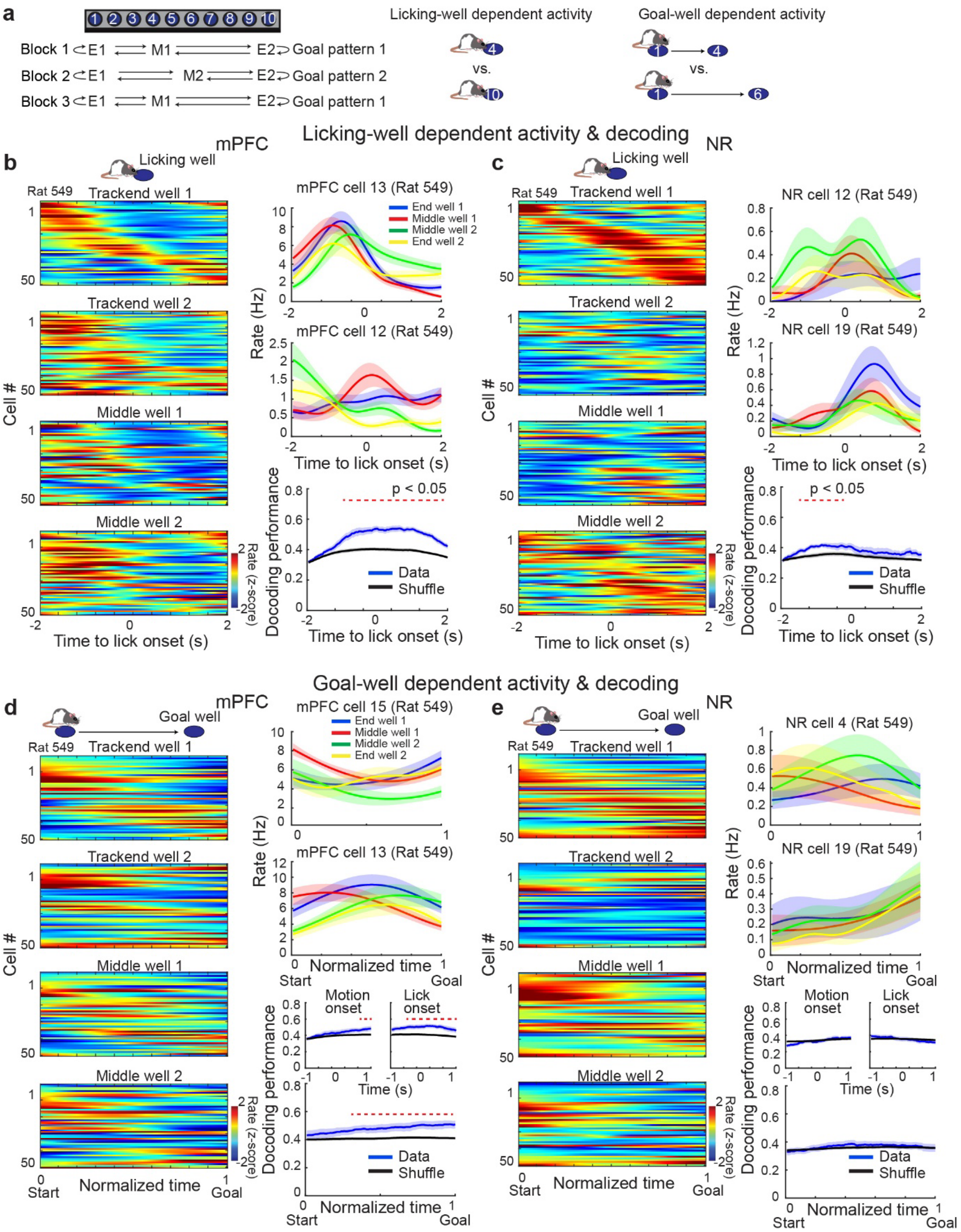


**Extended Data Figure 5: Representations of licking wells and goal wells in mPFC and NR.**

**a,** Schematic of the task structure, illustrating the distinction between licking-well-dependent and goal-well-dependent activity. **b,** Licking-well-dependent activity in representative mPFC neurons. Left, color-coded activity of 50 representative neurons aligned to well licking. Right, mean firing rates of two example neurons and decoding performance for licking-well identity. For decoding, one trial was used as the test set and the remaining trials as the training set (leave-one-out cross-validation). An LDA decoder was trained on z-scored firing-rate vectors in a window from −2 s to 2 s relative to lick onset, with trial numbers equalized across classes. Decoder performance was evaluated on the held-out test trial and repeated for all trials. Chance levels were estimated using two approaches: (i) training the decoder on data with shuffled class labels, and (ii) controlling for running direction by splitting trials by direction and shuffling class labels within each direction group, providing an estimate of decoding based solely on directional differences. The higher of the two chance levels was used. Red dashed lines indicate $p < 0.05$ in the Wilcoxon rank-sum test. **c,** Same as **b** for NR neurons. **d,** Goal-well–dependent activity in representative mPFC neurons. Left, color-coded activity of 50 representative neurons during navigation. Right, mean firing rates of two example neurons and decoding performance for goal-well identity. An LDA decoder was trained on z-scored firing-rate vectors in windows from −1 s to navigation onset and from −1 s to goal-well licking onset. Chance level estimation was identical to that in **b**. Red dashed lines indicate $p < 0.05$ in the Wilcoxon rank-sum test. **e,** Same as **d** for NR neurons. These results together suggest that information about individual goal locations is unlikely to be transmitted through the mPFC–NR–CA1 circuit.

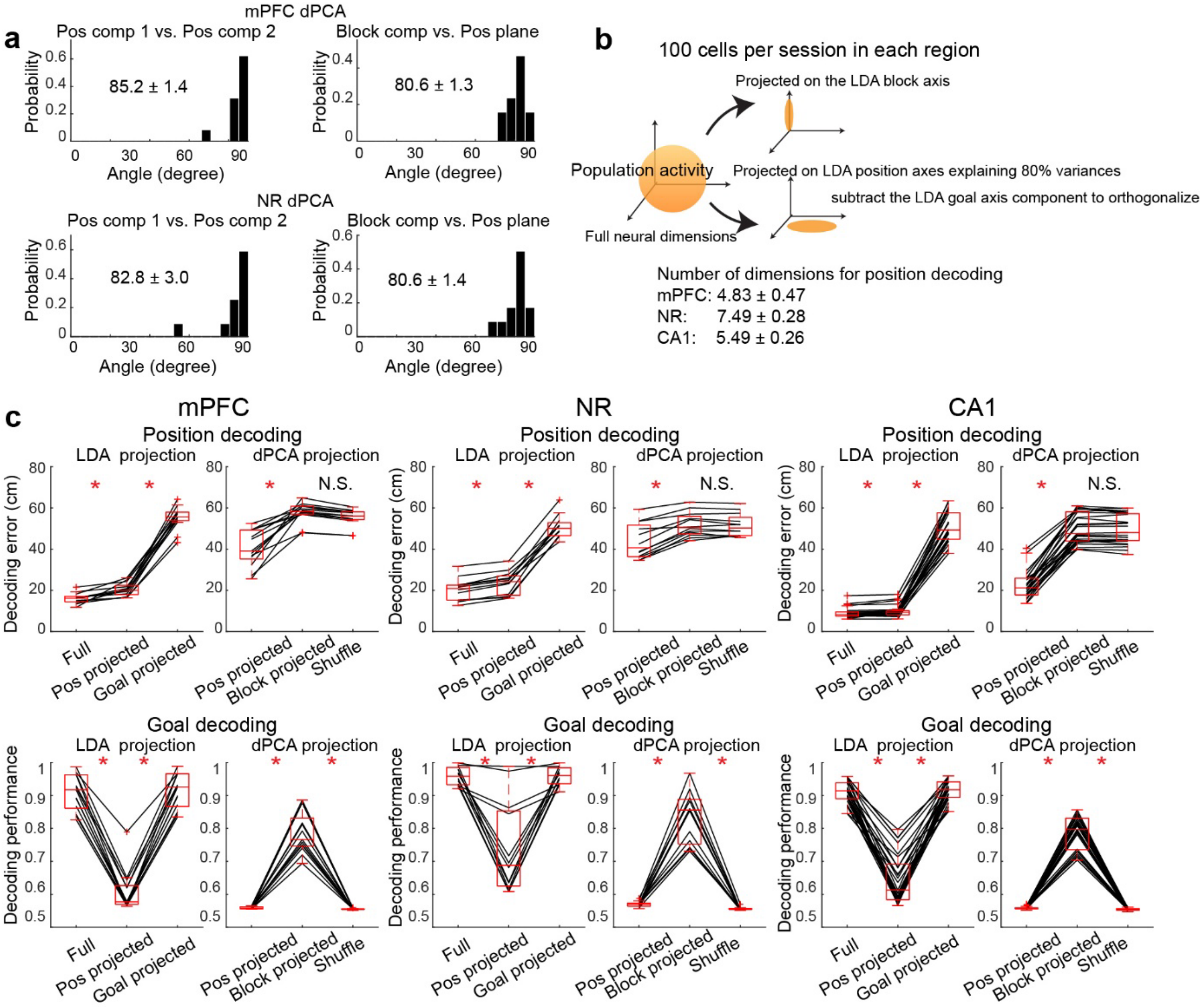


**Extended Data Figure 6: Performance comparison of goal-state and position coding across mPFC, NR, and CA1.**

**a,** Angles between the two position components and between the block component and the two position components extracted by dPCA, demonstrating near-orthogonality of the axes. **b,** As in Extended Data Fig. 3i, a goal classifier was constructed using linear discriminant analysis (LDA) to discriminate the two goal patterns, yielding a single projection axis corresponding to the goal-coding dimension. An LDA decoder for spatial position was also constructed, yielding 39 (40 − 1) projection axes. Dimensionality was reduced by retaining components explaining 80% of the variance. From this position-coding subspace, the component projected onto the goal-coding axis was subtracted, yielding a position subspace orthogonal to the goal-coding axis. An equal number of neurons (100 cells) was used for each brain region. Spatial and goal information were then compared using LDA decoding based on the full neural space, the orthogonalized position-coding subspace, or the goal-coding axis alone. **c,** Summary of decoding performance across brain regions for the LDA-based projections defined in **b** and for dPCA-derived projections.

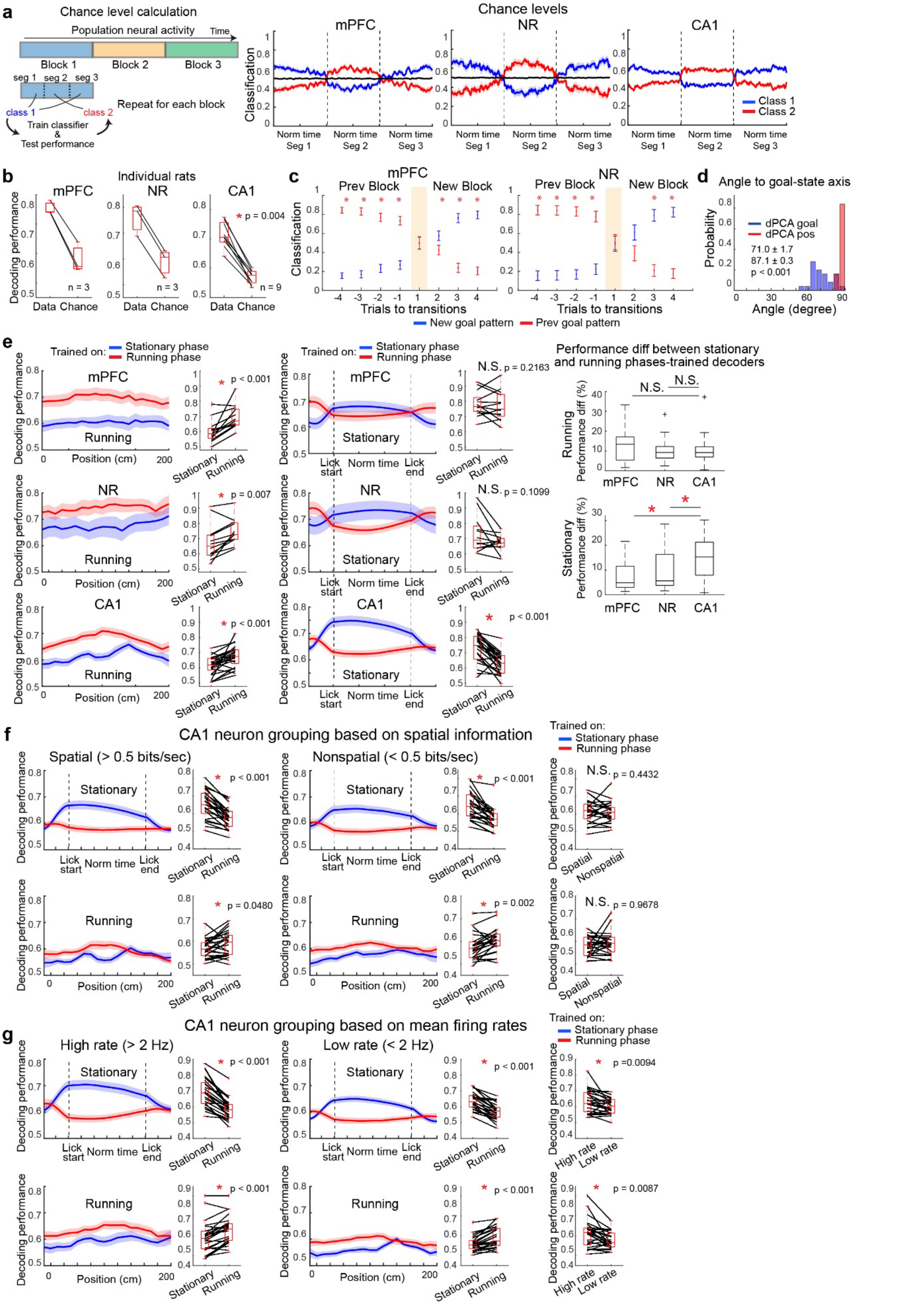


**Extended Data Figure 7: Chance-level estimation and locomotor-state independence of goal-state coding in mPFC, NR, and CA1.**

**a,** Estimation of chance-level goal-pattern decoding performance. To control for decoder performance unrelated to goal changes, three consecutive segments within the same goal block were artificially assigned to different classes (class 1 for the first and third segments, class 2 for the second segment), and the same decoding procedure was applied. This analysis was repeated independently for each of the three blocks, and decoding performance was averaged across blocks to estimate chance-level performance. The three plots on the right show the resulting chance levels for each brain region. **b,** Box plots showing goal-pattern decoding performance for individual rats in each brain region, demonstrating consistently higher performance than the chance levels estimated in **a**. **c,** Trial-by-trial goal-pattern decoding performance aligned to block transitions. Decoding performance is significantly above chance ($p < 0.05$ in Wilcoxon signed-rank test) and emerges rapidly following block transitions. During the first cycle after each transition (shaded), water was pre-filled at the goal wells and LEDs beneath the maze were illuminated to facilitate recognition of the new goal configuration. **d,** Comparison of angles between the goal-state classification axis (as in Fig. 4a) and the dPCA-derived axes. The goal-state axis is nearly orthogonal to the dPCA position components and more closely aligned with the dPCA block component, though not identical, likely reflecting differences between locomotor and immobility states used in the respective analyses. **e,** State dependence of goal-state representations. Goal-state classifiers were trained using neural activity from either stationary or running phases and then applied to both behavioral states. Box plots summarize decoding performance. During running, classifiers trained on running-phase data outperformed those trained on stationary data. In contrast, during stationary phases, classifiers trained on running-phase data performed comparably to those trained on stationary data in mPFC and NR, but not in CA1. This asymmetry indicates that goal-related information encoded during running generalizes to stationary states in mPFC and NR, but not in CA1, consistent with a role for mPFC–NR inputs in stabilizing CA1 goal-state coding across transitions from locomotion to immobility. $p < 0.05$ in Wilcoxon rank-sum test. **f,** Contribution of CA1 cell types to goal-state decoding. CA1 neurons were divided into two groups based on spatial information content (high or low relative to 0.5 bits/s). Cell numbers were matched between groups within each session before decoding analysis. No significant difference in decoding performance was observed, indicating that goal-state coding is not preferentially driven by either spatially tuned or weakly spatial neurons. **g,** Contribution of CA1 neurons categorized by mean firing rate. CA1 neurons were divided into high- and low-firing-rate groups (threshold: 2 Hz), with cell numbers matched across groups. Goal-pattern decoding performance was significantly higher when using high-firing-rate neurons, suggesting that neurons with higher firing rates contribute disproportionately to goal-state coding.

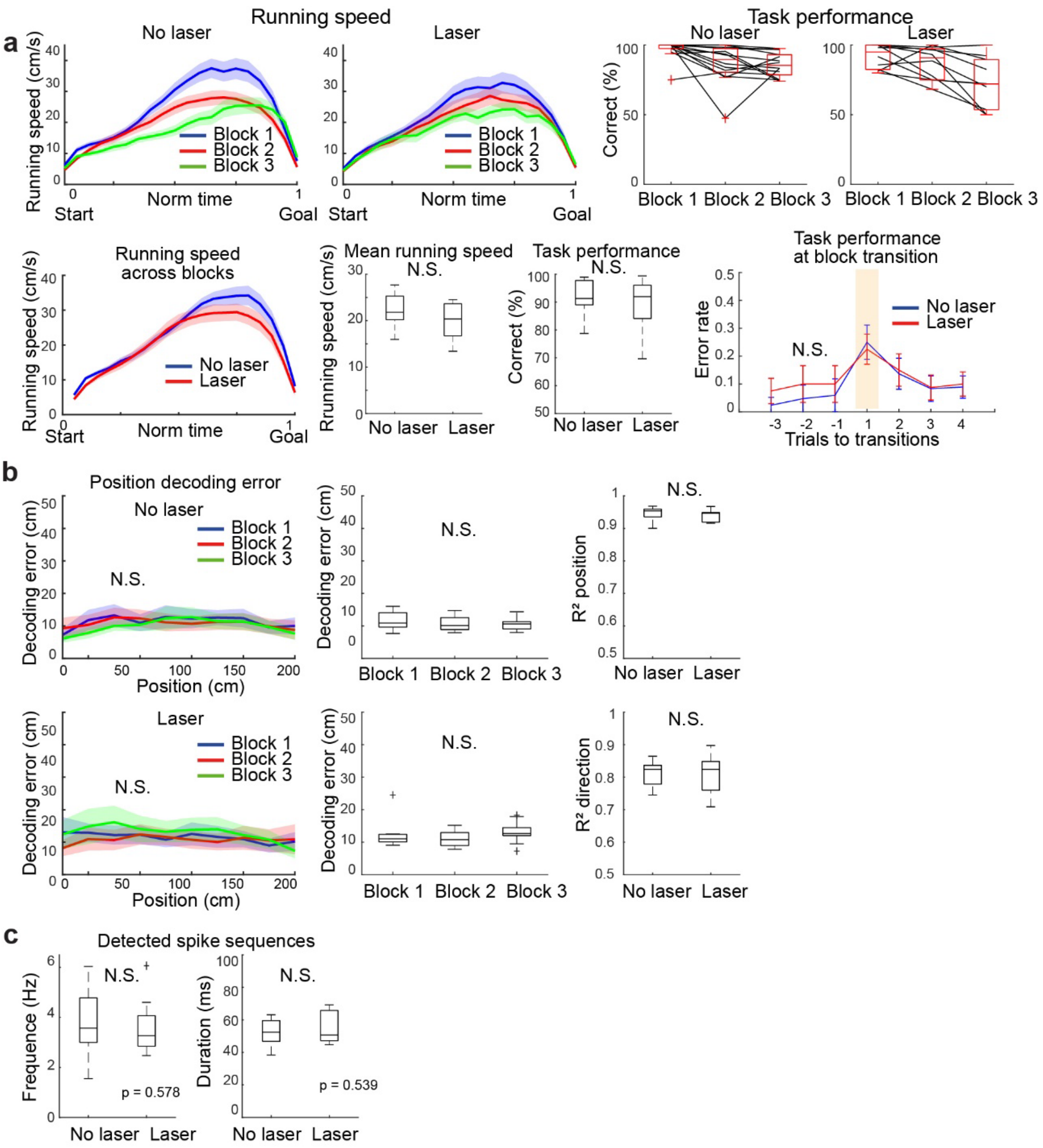


**Extended Data Figure 8: Non-significant effects of NR silencing on task performance, position decoding, and CA1 spike sequence duration and frequency.**

**a,** Top left: Running speed during goal-directed journeys across blocks with and without laser application. Top right: Box plots showing task performance across blocks with and without laser. Bottom left: Comparison of average running speed between laser and no-laser sessions. Bottom middle: Box plots comparing running speed and task performance. No significant differences were observed between conditions. Bottom right: Error rates during block transitions, showing no significant difference between laser and no-laser sessions. $p > 0.05$, Wilcoxon rank-sum test. **b,** Left: Position-decoding errors from CA1 population activity across maze positions with and without laser application. Middle: Comparison of position-decoding errors across blocks, showing no significant differences. $p > 0.05$, Kruskal–Wallis test. Right: Box plots comparing the coefficient of determination ($R^2$) for decoded position and running direction between laser and no-laser sessions, showing no significant differences. $p > 0.05$, Wilcoxon rank-sum test. **c,** Comparison of detected spike-sequence events. Both the frequency and duration of detected events show no significant differences between laser and no-laser sessions. $p > 0.05$, Wilcoxon rank-sum test.

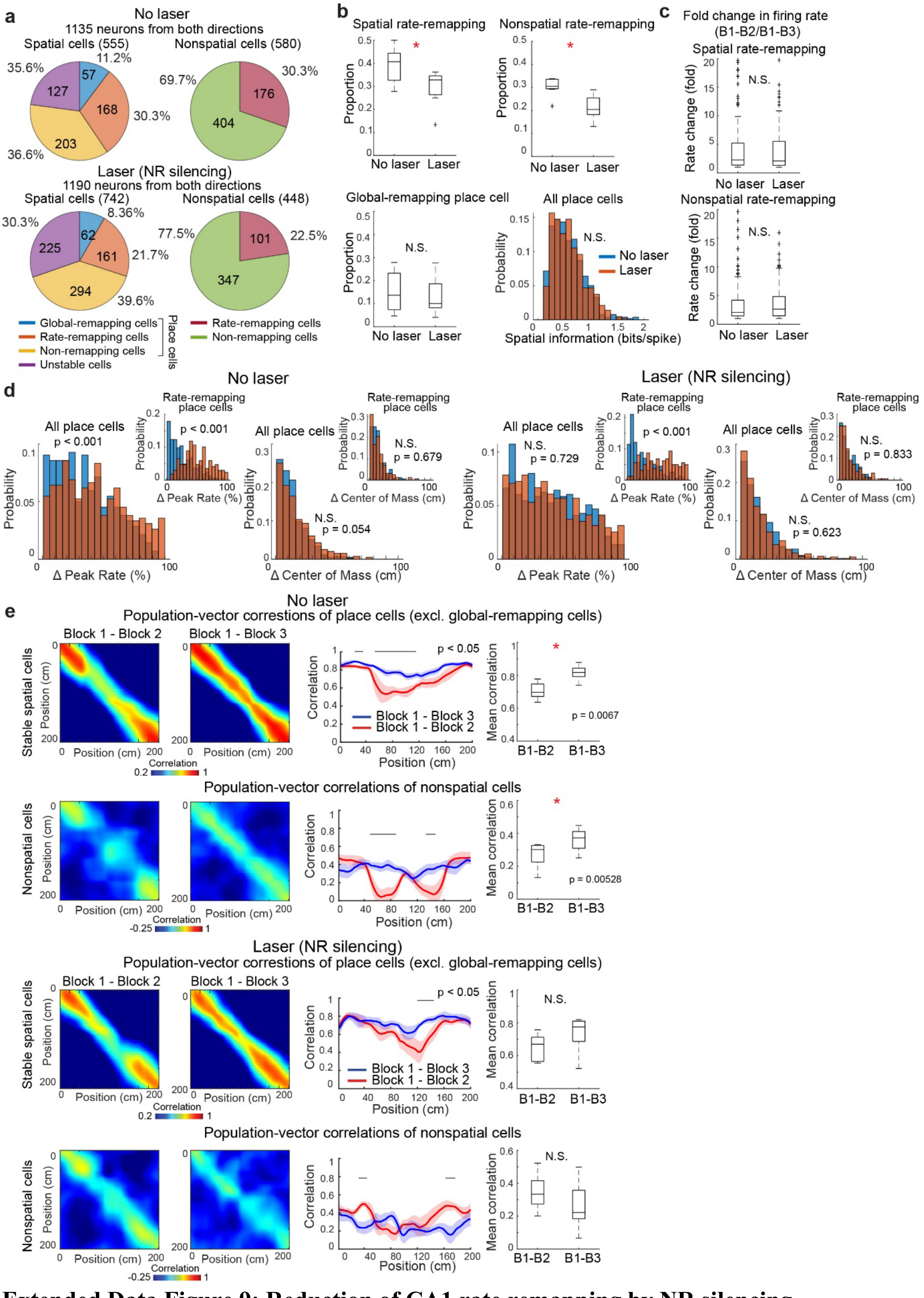


**Extended Data Figure 9: Reduction of CA1 rate remapping by NR silencing.**

**a,** Pie charts showing the proportions of different CA1 cell categories, plotted separately for spatial and nonspatial neurons. Data are shown for 4 animals, with sessions performed with and without laser application (8 sessions per condition). **b,** Comparison of the proportions of rate-remapping place cells (top left), nonspatial rate-remapping cells (top right), and global-remapping place cells (bottom left). Significant differences were observed in the proportion of rate-remapping cells between sessions with and without laser application (rate-remapping place cells: Friedman test, $\chi^2(1) = 5.40$, $p = 0.020$; nonspatial rate-remapping cells: $\chi^2(1) = 5.40$, $p = 0.020$) but not in global-remapping cells: $\chi^2(1) = 0.60$, $p = 0.439$). Bottom right: Distribution of spatial information for all place cells, showing no significant difference between laser and no-laser sessions (Kolmogorov–Smirnov test, $p = 0.934$). **c,** Fold changes in firing-rate modulation comparing blocks 1–2 (different goal patterns) with blocks 1–3 (same goal pattern) within rate-remapping place cells (top) and nonspatial rate-remapping cells (bottom). No significant differences were observed between laser and no-laser sessions. $p > 0.05$, Wilcoxon rank-sum test. **d,** Histograms showing distributions of peak firing rate differences between the same goal blocks (blue) and the different goal blocks (red), comparing sessions with and without laser application. In control sessions, place cells exhibited a significant difference in the distribution of rate changes between blocks 1–2 and 1–3 (*$p < 0.001$, Kolmogorov–Smirnov test), whereas this difference was abolished during NR silencing ($p = 0.729$). Center-of-mass shifts of place fields remained stable across blocks with different goals and were unaffected by NR silencing. **e,** Same analysis as in Extended Data Fig. 3g, comparing sessions with and without laser application. Black lines in the middle panels indicate significant effects ($p < 0.05$, Friedman test). Right: Comparison between blocks 1–2 and 1–3. Mean population-vector (PV) correlations for place cells excluding global-remapping cells were significantly different in control sessions ($\chi^2(1) = 7.35$, $p = 0.007$) but not during NR silencing ($\chi^2(1) = 3.75$, $p = 0.053$). For nonspatial rate-remapping cells, significant differences were observed without laser ($\chi^2(1) = 9.60$, $p = 0.002$) but not with laser ($\chi^2(1) = 0.00$, $p = 1.000$). Data are from 8 sessions across 4 animals.

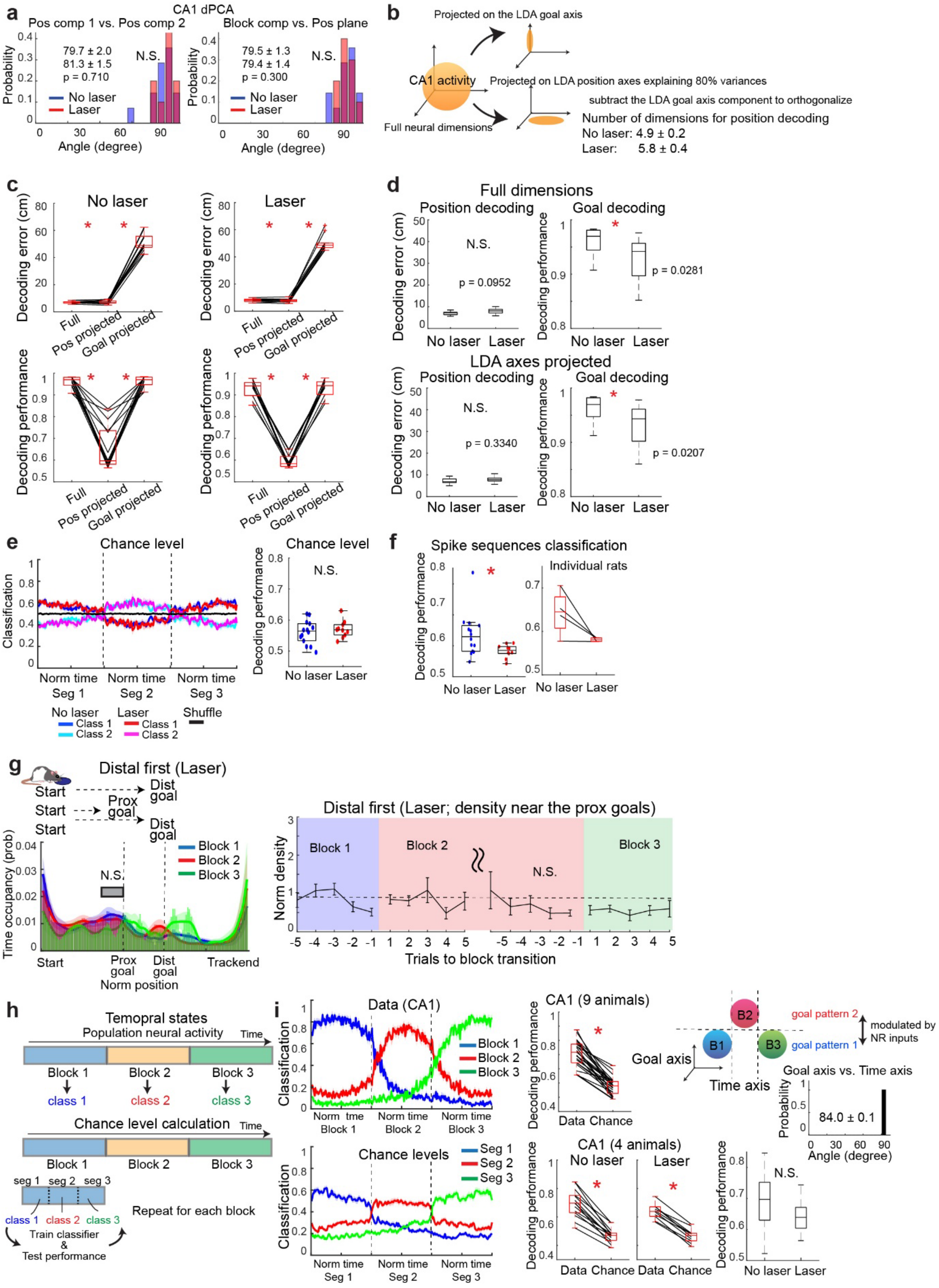


**Extended Data Figure 10: Selective effects of NR silencing on CA1 goal-state coding and pre-navigation spike sequences.**

**a,** Comparison of angles between dPCA-derived components in sessions with and without laser application. No significant difference was observed. p > 0.05, Kolmogorov–Smirnov test. **b,** As in Extended Data Fig. 3i, a goal-pattern classifier was constructed using linear discriminant analysis (LDA) to discriminate between the two goal patterns, yielding a single projection axis corresponding to the goal-coding dimension. An LDA decoder for spatial position was also constructed, producing 39 (40 − 1) projection axes. Dimensionality was reduced by retaining components explaining 80% of the variance. The projection of the goal-coding axis was then subtracted from the position-coding subspace, yielding a position subspace orthogonal to the goal-coding dimension. **c,** Summary of decoding performance for sessions with and without laser application using the LDA-based projections defined in **b**, shown as position decoding error (top) and goal-pattern classification accuracy (bottom). No significant differences were detected. $p > 0.05$, Wilcoxon signed-rank test. **d,** Comparison of decoding performance using full-dimensional LDA classifiers (top) and LDA-based orthogonal projections defined in **b** (bottom). $p < 0.05$, Wilcoxon rank-sum test. **e,** Estimation of chance-level goal-pattern decoding performance, as in Extended Data Fig. 7a. Chance-level performance did not differ between sessions with and without laser application. **f,** Classification performance of pre-navigation spike sequences in sessions with and without laser application, shown for all sessions (left) and for individual animals (right). $p < 0.05$, Wilcoxon rank-sum test. **g,** Same analysis as in Fig. 1f applied to sessions with nucleus reuniens (NR) silencing. Left, time-occupancy probabilities of pre-navigation spike sequences for trials targeting a distal goal in blocks 1 and 3 and a proximal goal in block 2. Right, trial-wise densities normalized relative to block 1. No significant differences were observed across blocks. $p > 0.05$, Wilcoxon signed-rank test. **h,** Classification performance of task-block progression (blocks 1–3) based on randomly sampled 2-s neural activity segments, irrespective of locomotor state (top), and the corresponding estimation of chance-level performance (bottom). **i,** Right, classification performance for temporal sequences of task blocks (top) and corresponding chance levels (bottom). Middle and right, box plots summarizing decoding performance across all sessions from nine animals (top) and comparisons of the same four animals with and without laser application (bottom). $p < 0.05$, Wilcoxon signed-rank test. The schematic (right top) illustrates orthogonal neural coding of goal-state and temporal sequence information, supported by the near-90° distribution of angles between these axes.
